# Monitoring Dynamics of Large Membrane Proteins by ^19^F Paramagnetic Longitudinal Relaxation: Domain Movement in a Glutamate Transporter Homolog

**DOI:** 10.1101/832121

**Authors:** Yun Huang, Xiaoyu Wang, Guohua Lv, Asghar M. Razavi, Gerard H. M. Huysmans, Harel Weinstein, Clay Bracken, David Eliezer, Olga Boudker

**Affiliations:** Department of Physiology & Biophysics, Weill Cornell Medicine, 1300 York Ave, New York, NY 10021; Department of Biochemistry, Weill Cornell Medicine, 1300 York Ave, New York, NY 10021; Howard Hughes Medical Institute, Chevy Chase

## Abstract

In proteins where conformational changes are functionally important, the number of accessible states and their dynamics are often difficult to establish. Here we describe a novel ^19^F-NMR spectroscopy approach to probe dynamics of large membrane proteins. We labeled a glutamate transporter homologue with a ^19^F probe via cysteine chemistry and with a Ni^2+^ ion via chelation by a di-histidine motif. We used distance-dependent enhancement of the longitudinal relaxation of ^19^F nuclei by the paramagnetic metal to assign the observed resonances. We identified two outward- and one inward-facing states of the transporter, in which the substrate-binding site is near the extracellular and intracellular solutions, respectively. We then resolved the structure of the unanticipated second outward-facing state by Cryo-EM. Finally, we showed that the rates of the conformational exchange are accessible from measurements of the metal-enhanced longitudinal relaxation of ^19^F nuclei.

## Introduction

The advent of crystallography and single-particle Cryo-EM led to a rapid expansion of protein structure determinations. Often, multiple conformations of the same protein are reported providing snapshots of functional states, but their temporal relationship is usually unknown. NMR spectroscopy is a versatile approach to probe protein dynamics ^1^, but measurements on large assemblies, such as membrane proteins, are challenging because of the severe broadening and overlap of ^1^H, ^15^N and ^13^C resonances. Recently developed TROSY NMR addresses the issue ^2,3^, but requires protein perdeuteration, which is cumbersome and not always possible for membrane proteins ^4^.

^19^F-NMR is a useful alternative because of its high sensitivity, robust chemical shift dispersion, the absence of background signals, and no requirement for perdeuteration ^5 6^. Fluorinated probes have been introduced into proteins via incorporation of fluorinated or unnatural amino acids ^7^ or via cysteine chemistry ^8^. Many methods to follow dynamics, including saturation transfer, EXSY, CPMG, CEST and DEST, have been adapted to ^19^F-NMR ^6,9-12^. One existing challenge, however, is assigning observed resonances to known structural states. Typically, the assignment relies on population shifts expected in response to ligand binding or mutations and, therefore, requires prior information from other techniques ^9^. Alternatively, differences in solvent paramagnetic resonance enhancement (PRE) ^8^ have been used, but this approach only works when the conformational states feature distinct solvent exposure of the ^19^F probe. Here, we undertook measurements of ^19^F longitudinal relaxation rates (*R*_1_) and their distance-dependent enhancement by paramagnetic ions chelated by di-histidine motifs to assign resonances to different states and to estimate the rates of the conformational exchange.

To develop the strategy, we used a membrane transporter, GltPh, which forms ∼300 kDa particles in detergent micelles ^13^. GltPh is an aspartate/sodium symporter from *Pyrococcus horikoshii*, homologous to human glutamate transporters, which play important roles in glutamatergic neurotransmission in the brain ^14^. Human glutamate transporters and GltPh utilize trans-membrane ionic gradients, primarily of sodium (Na^+^) ions, to drive concentrative uptake of their substrates ^15,16^. GltPh transports its cargo across the membrane by isomerizing between outward-facing states (OFS), where the substrate and ion binding sites are accessible from the extracellular solution, and inward-facing states (IFS) where they are accessible from the cytoplasm (**Fig. 1a**). The dynamics of these conformational transitions are thought to determine the rate of transport ^17^. GltPh is a homotrimeric protein in which each subunit functions independently ^18^. The transition between the OFS and the IFS involves a concerted ∼15 Å transmembrane movement of a peripherally located transport domain, which harbors the substrate- and ion-binding sites, relative to the largely stationary central trimerization scaffold domain. We labeled a cysteine introduced into the transport domain of GltPh with a ^19^F trifluoroethanethio (TET) moiety ^19^ (**Fig. S1a**) and placed a pair of histidines into a transmembrane helix (TM) of the scaffold domain to chelate a nickel (Ni^2+^) ion at a site where the distance to the ^19^F probe changes upon state transition (**Fig. 1a and 1b**). By measuring ^19^F longitudinal relaxation rates and Ni^2+^-mediated PRE, we assigned the three resonances observed in the ^19^F spectrum of the transporter to one IFS and two distinct OFS conformations. Guided by the NMR experiments, we determine the Cryo-EM structures of the two outward-facing states. Finally, we used measurements of Ni^2+^-enhanced *R*_1_ relaxation to estimate the exchange rate between the OFS and the IFS conformations.

**Figure 1:**
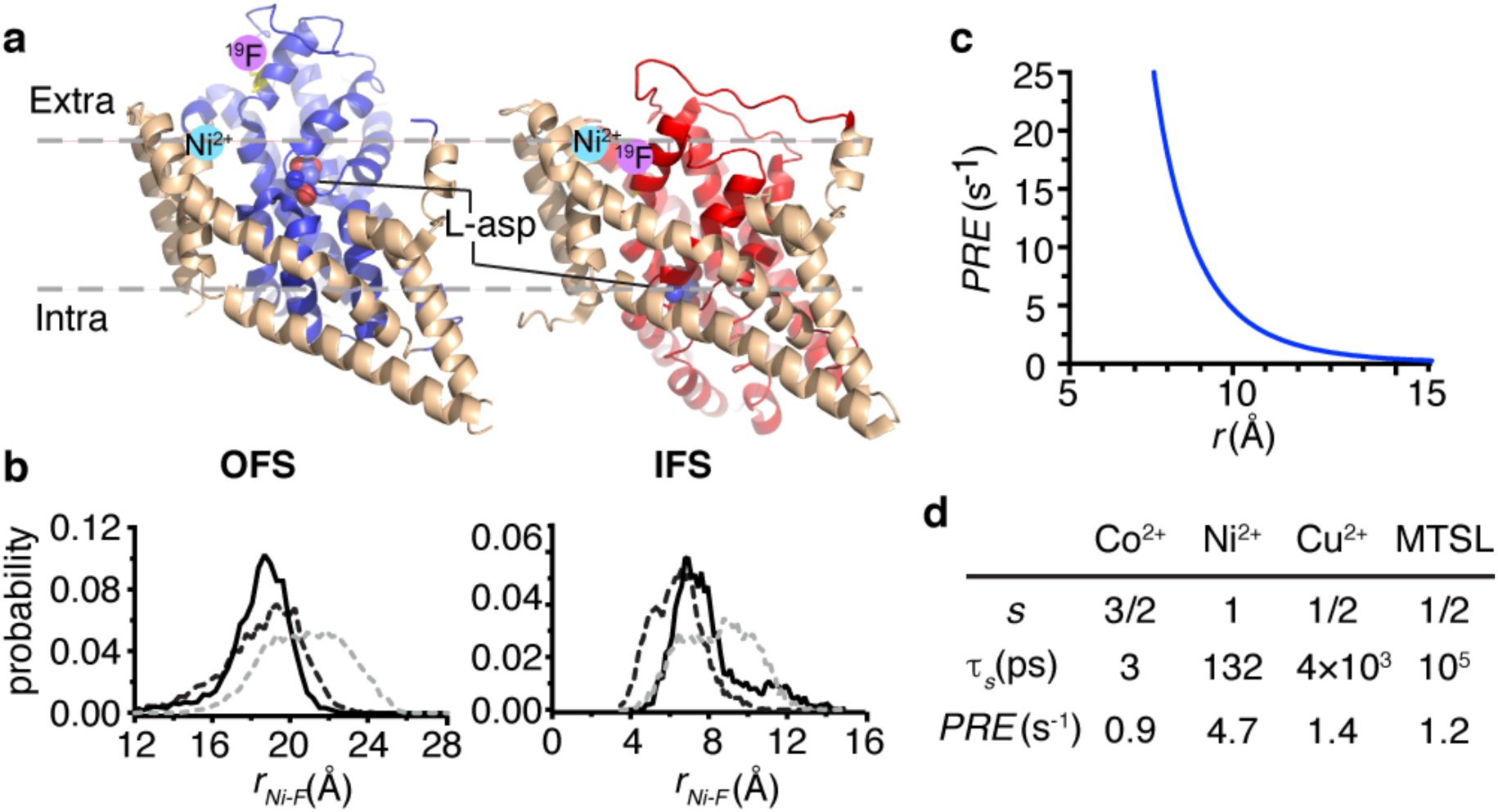
Design for ^19^F and Ni^2+^ labeling of GltPh for *R*_*1*_ *PRE*. **a**, Cartoon representations of the structures of L-asp bound GltPh in the OFS (PDB accession code 2NWX) and IFS (accession code 3KBC). The structurally rigid scaffold domain is colored wheat and the dynamic transport domain is colored blue and red in the OFS and IFS, respectively. The substrate L-asp is shown as spheres. Pink and cyan circles represent the expected locations of the ^19^F label and bound Ni^2+^ ion, respectively. **b**, Distance probability distributions between ^19^F and Ni^2+^ calculated from 100 ns of the molecular dynamics simulation trajectories. Black solid, black dashed and gray dashed lines correspond to protomers A, B and C, respectively. **c**, Dependence of the longitudinal *R*_*1*_ PRE on the distance between ^19^F and Ni^2+^ based on Equation 1. **d**, *R*_*1*_ *PRE* calculated for ^19^F nucleus at a distance of 10 Å from different paramagnetic centers, assuming τ_*c*_ is 213 ns, *S*^2^ is 0.1 and τ_*i*_ is 20 ps. *s*, spin quantum number. τ_*s*_, electron relaxation time ^49-52^. MTSL, (1-Oxyl-2,2,5,5-tetramethylpyrroline-3-methyl)methanethiosulfonate.

## Results

### Expected Ni^2+^-mediated ^19^F longitudinal relaxation enhancement

PRE results from a dipole-dipole interaction between spins of an unpaired electron and a nucleus, which depends on the inverse sixth power of the distance, *r,* between them. The transition metal ions Mn^2+^, Cu^2+^, Ni^2+^ and Co^2+^ and organic nitroxide radicals have been used as paramagnetic centers in EPR ^20^, solution NMR ^21^ and solid state NMR measurements ^22-25^. We aim to measure the magnitude of the PRE effect on the longitudinal relaxation, *R*_1_ PRE, which is described by the Solomon-Bloembergen equations ^26,27^:

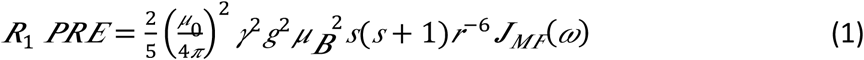

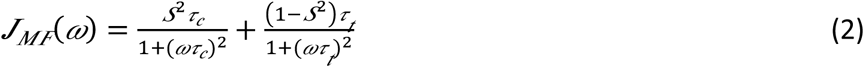

where *μ*_*0*_ is the permeability of vacuum, *γ* is the nuclear gyromagnetic ratio, *g* is the electron g-factor, *μ*_*B*_ is the magnetic moment of the free electron, *s* is the electron spin quantum number, *ω* is the nuclear Larmor frequency and *S*^2^ is the order parameter. *J*_*MF*_ is the spectral density function accounting for the local side chain motions (**Supplementary Information**) ^28,29^.

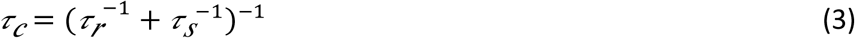

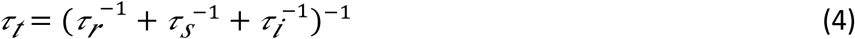

where τ_*r*_ is the protein rotation correlation time, τ_*s*_ is the electron relaxation time, and τ_*i*_ is the characteristic internal motion time.

Because PRE depends quadratically on the gyromagnetic ratio, it is weak for ^13^C and ^15^N nuclei but is significant for ^1^H and ^19^F ^30^. Typically, the enhancement of the transverse relaxation rate (*R*_2_ *PRE*) of ^1^H nuclei is used to obtain distance information. The approach is widely used to facilitate structure determination, identify protein-protein interfaces and detect invisible transient states ^31^. In contrast, retrieving the paramagnetic component from the enhanced longitudinal relaxation rate (*R*_1_ *PRE*) for ^1^H nuclei is difficult because of the interference from ^1^H-^1^H cross relaxation ^31^. The transverse relaxation time of ^19^F nuclei in membrane proteins is very short, only a few milliseconds ^8^, rendering the *R*_2_ *PRE* effect insignificant. However, ^19^F longitudinal relaxation in proteins occurs on a seconds time scale and is dominated by dipole-dipole interactions ^32^ with negligible heteronuclear ^19^F-^1^H cross relaxation ^33,34^. Notably, because of the negligible contributions of ^1^H-^19^F cross relaxation, ^19^F longitudinal relaxation is effectively mono-exponential ^33^.

Paramagnetic Ni^2+^ ions can be chelated by pairs of histidines placed into *i* and *i*+4 positions of a helix ^35^, and the resultant PRE can be measured by taking the difference of the relaxation rates in the presence of an ion, *R*_*1, Ni*_ and in its absence, *R*_*1,ref*_:

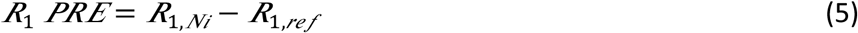

Based on Equation 1, we estimated the expected *R*_1_ *PRE* for a ^19^F nucleus at a distance of 10 Å from a paramagnetic center using a τ_*r*_ of 213 ns. Considering that a TET-modified cysteine side chain is similar in size to methionine, we used a value of 0.1 for the order parameter *S*^2^ and 20 ps for the internal motion time τ_*i*_, which are typical values found for methionine methyl groups ^36,37^. Ni^2+^ ion is advantageous compared to other paramagnetic centers because it has an intermediate electron relaxation time of 132 ps ^38^ that results in a larger enhancement of the nuclear longitudinal relaxation, *R*_1_ *PRE* ^39^ (**Fig. 1d**). As PRE drops off steeply with the distance between the unpaired electron and the ^19^F nucleus, we estimate the longest measurable distance to be ∼14 Å (**Fig. 1c**). Notably, due to the dominant contribution of τ_*i*_ and τ_*s*_ in Equations 3 and 4, the *R*_1_ *PRE* is not sensitive to the protein size.

### ^19^F-NMR spectra of GltPh

We set out to use ^19^F *R*_1_ *PRE* to detect and distinguish the two key functional states of GltPh, the OFS and the IFS (**Fig. 1a**). Therefore, we introduced a single cysteine mutation M385C into the transport domain for labeling with TET and a double histidine mutation Y215H/E219H (dHis) into TM5 of the scaffold domain to generate a coordination site for a Ni^2+^ ion (**Fig. 1a**). M385C GltPh and dHis/M385C GltPh proteins were purified, efficiently labeled with TET by a two-step method ^19^, and found to remain active in L-aspartate (L-asp) uptake experiments when reconstituted into proteoliposomes (**Supplementary Fig. S1**).

1D ^19^F-NMR spectra of M385C-TET GltPh (**Fig. 2a**) and dHis/M385C-TET GltPh (**Supplementary Fig. S2**) in the presence of saturating concentrations of Na^+^ ions and L-asp were indistinguishable from each other and showed three partially overlapping resonances, S1-3. Notably, dHis/M385C-TET GltPh in complex with a non-transportable blocker DL-*threo*-3-methyl aspartic acid (TMA) ^40^ displayed a similar spectrum (**Fig. 2d)**. Thus, the ^19^F-NMR probe revealed the presence of three distinct conformations of the transporter that did not undergo rapid exchange on the NMR time scale. Indeed, slow interconversion between the OFS and the IFS on the order of seconds is expected for GltPh based on single-molecule FRET (smFRET) and atomic force microscopy (AFM) studies ^17,41,42^. Earlier measurements also showed that Na^+^ and L-asp-bound GltPh samples the OFS and the IFS with comparable probabilities ^17,43^. Therefore, it is possible that S1, S2, and S3 peaks, which all show similar populations, correspond to either OFS or IFS. To assign the resonances to specific states, we first examined how distinct ligands changed their populations. In the absence of L-asp, GltPh binds Na^+^ with about 3-fold higher affinity in the OFS than in the IFS, suggesting that Na^+^ alone should favor the OFS ^44^. The ^19^F-NMR spectrum of dHis/M385C-TET GltPh in the presence of 0.6 M Na^+^ and in the absence of L-asp showed a diminished S1 peak (**Fig. 2a** and **b**), suggesting that the peak corresponded to the IFS. Consistently, the S1 peak also disappeared in the presence of a competitive blocker DL-threo-β-benzyloxyaspartic acid (TBOA) (**Fig. 2c**), which favors the OFS ^45^. The S2 and S3 peaks persisted under these conditions, suggesting that both corresponded to the OFS. If so, the relative populations of the IFS and OFS of Na^+^/L-asp-bound dHis/M385C-TET GltPh calculated from the areas of the 3 peaks, were 30 ± 1.4 % and 70 ± 6 %, respectively, consistent with previous measurements ^17,43^. To support our peak assignments, we collected 1D ^19^F spectra for dHis/M385C-TET GltPh bearing additional mutations that favor the IFS, K290A and R276S/M395R (termed RSMR from here on, for brevity) ^17,41^. Both mutants showed similar peak positions to dHis/M385C-TET GltPh, but dramatically increased intensities of peak S1, consistent with this resonance corresponding to the IFS (**Fig. 2e,f**).

**Figure 2:**
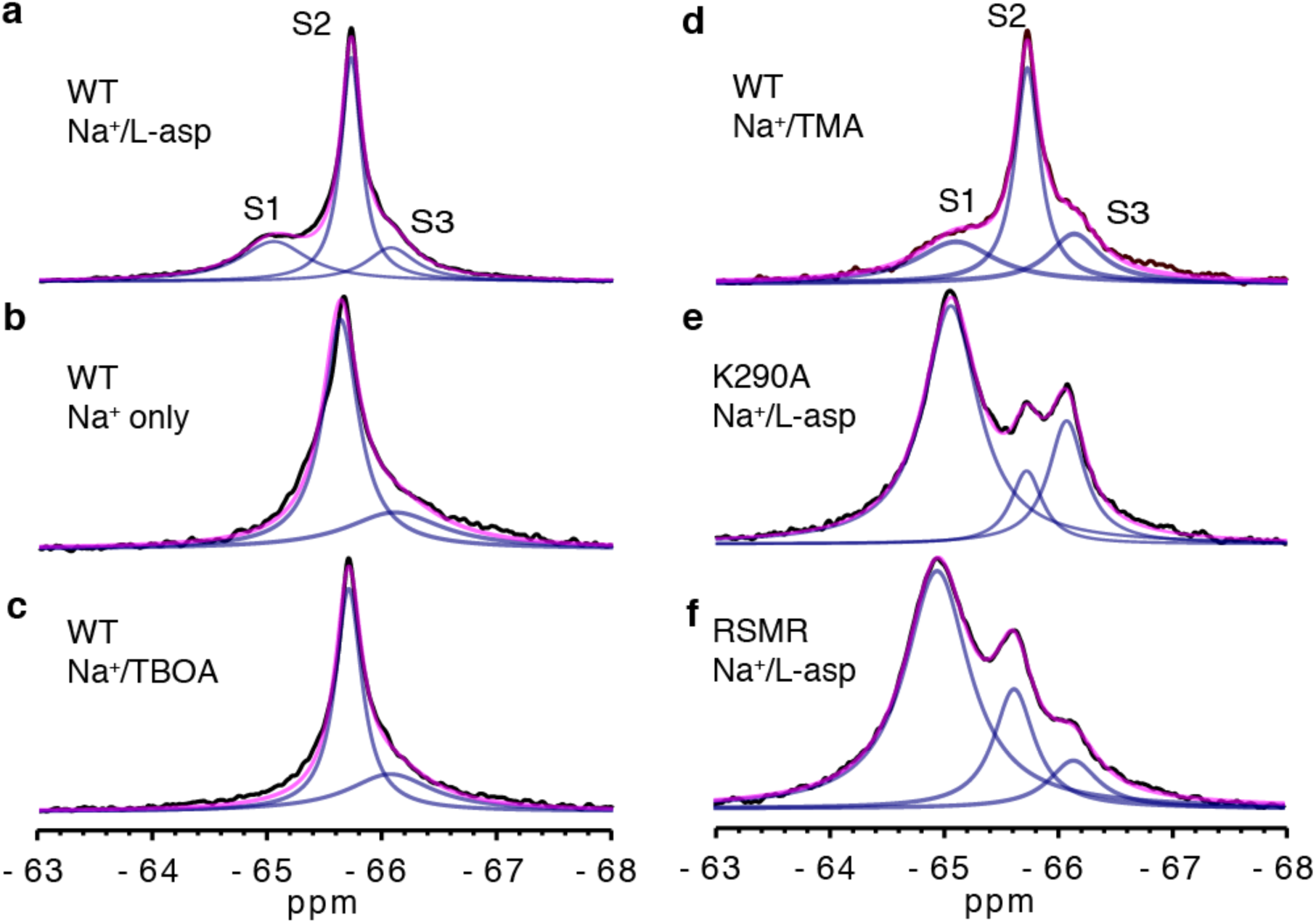
^19^F-NMR spectra of dHis/M385C-TET GltPh and its mutants. 1D spectra of dHis/M385C-TET GltPh (WT) recorded at 293 K in the presence of 200 mM Na^+^ and 10 µM L-asp (**a**), 0.6 M Na^+^ only (**b**), 200 mM Na^+^ and 1 mM TBOA (**c**) or 200 mM Na^+^ and 110 µM TMA (**d**). 1D spectra of K290A dHis/M385C-TET (**e**) and RSMR dHis/M385C-TET (**f**) GltPh mutants in the presence of 200 mM Na^+^ and 10 µM L-asp. The spectra were deconvoluted into Lorentzian peaks S1, S2 and S3. Raw data are black, fits are pink and deconvoluted peaks are light blue.

### Ni^2+^-mediated PRE identifies two outward- and one inward-facing states

To measure Ni^2+^-mediated PRE, we added 3 molar equivalents of NiSO_4_ to dHis/M385C-TET GltPh. Unexpectedly, the S1 peak showed increased intensity, corresponding to a population increase to 60 ± 4.5 % (**Fig. 2a, 3a and Supplementary Fig. S2**). In contrast, the control M385C-TET GltPh lacking the dHis motif did not show this population change (**Supplementary Fig. S2**). Thus, it appears that Ni^2+^ binding to the dHis motif in TM5 favors the IFS by ∼1.3 kT. The dHis motif is located at the interface between the transport and scaffold domains, and the local environment around the motif may differ in the OFS and the IFS, leading to higher affinity for Ni^2+^ in the latter. Titrating dHis/M385C-TET GltPh with Ni^2+^ and plotting the intensity of the S1 peak yielded a binding isotherm (**Fig. 3a**) from which we estimated the affinity for the Ni^2+^ ion of 34 ± 16 μM. This affinity is within the range of values reported for transition metal ions binding to dHis motifs ^46^.

**Figure 3:**
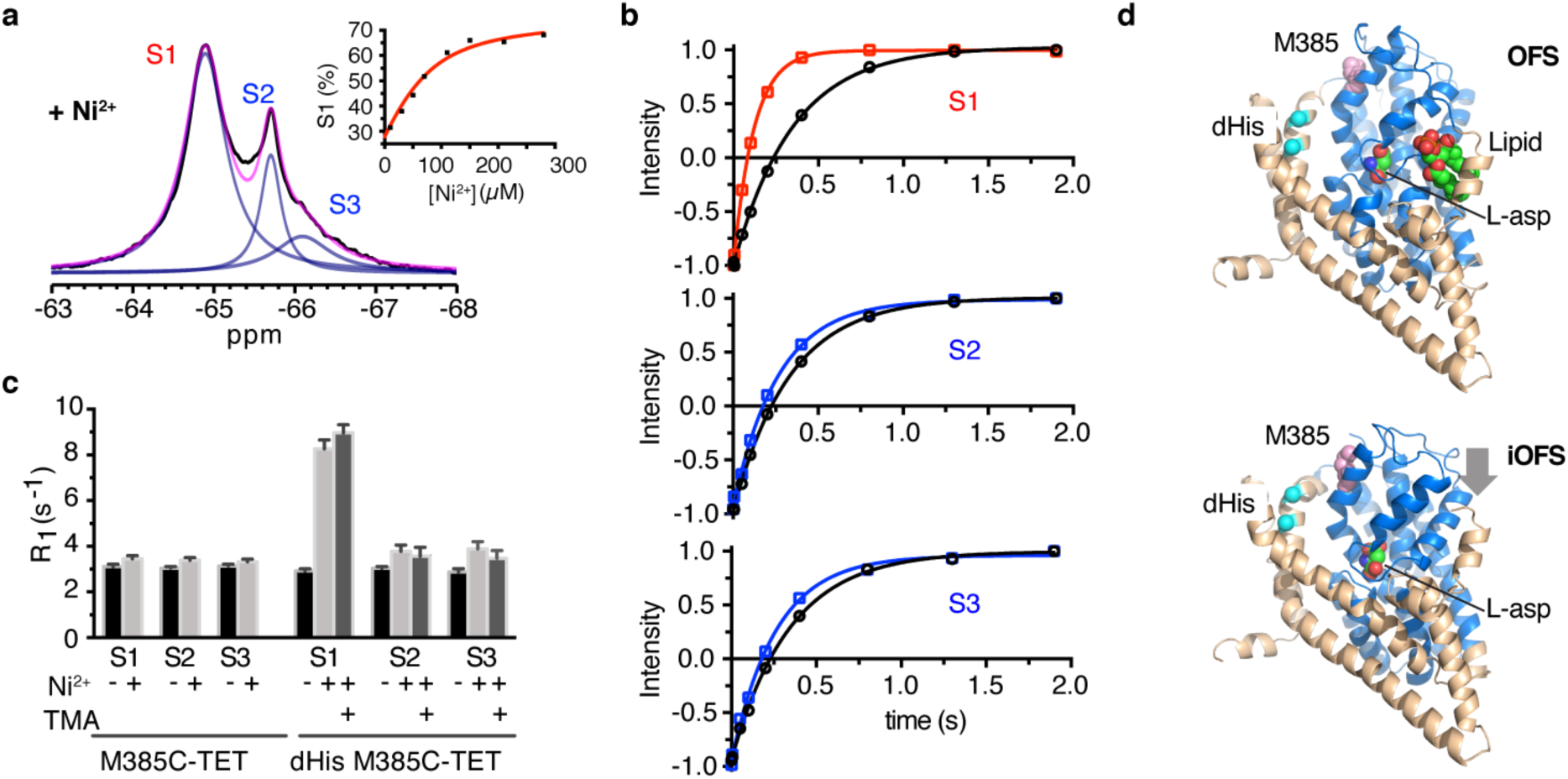
^19^F peak assignment using Ni^2+^-mediated PRE. **a**, 1D ^19^F spectra of Na^+^/L-asp-bound dHis/M385C-TET GltPh recorded at 293 K in the presence of 3 molar equivalents of Ni^2+^ ions. Inset: Ni^2+^ titration of 69 μM protein. Data are plotted as peak S1 population. Solid red line is the fit to the quadratic binding equation. **b**, *R*_*1*_ relaxation measured for S1 (top), S2 (middle) and S3 peak (bottom) in the absence (black) and in the presence of Ni^2+^ ions (color). Solid lines through the data correspond to mono-exponential fits and the results of the fits are shown in **Supplementary Table S1. c**, *R*_*1*_ rates of S1, S2 and S3 peaks of M385C-TET and dHis/M385C-TET GltPh constructs in the absence (black) and presence of Ni^2+^ ions and bound with L-asp (silver) or TMA (gray). All samples contained NaCl and L-asp, except where indicated L-asp was replaced with TMA. **d**, Cartoon representation of the Cryo-EM structures of the wild type GltPh, which showed the OFS (top) and the iOFS (bottom) conformations. The scaffold and transport domains are colored wheat and blue, respectively. Bound substrate L-asp and a lipid model are shown as spheres and colored by atom type. M385 is rendered as pink spheres. The location of the dHis motif is highlighted as cyan spheres. The broad grey arrow near the in the iOFS model indicates an inward motion of ∼5 Å of the transport domain.

Next, we measured *R*_1_ relaxation rates for each peak in the absence and in the presence of Ni^2+^ ions. We observed a relatively strong PRE of 5.4 ± 0.4 s^−1^ for peak S1, while peaks S2 and S3 showed weak PRE effects of 0.7 ± 0.3 s^−1^ and 1.0 ± 0.3 s^−1^, respectively (**Fig. 3b** and **c**). When L-asp was replaced by the non-transportable blocker TMA, we observed slightly higher PRE values for peak S1 and slightly lower ones for peaks S2 and S3 (**Fig. 3c** and **Supplementary Table S1**). In contrast, M385C-TET GltPh without the dHis motif did not show significant *R*_1_ *PRE* in the presence of Ni^2+^ ions for either of the peaks (**Fig. 3c**). Using Equation 1 and PRE values measured for dHis/M385C-TET GltPh in the presence of TMA, we estimated Ni^2+^ to ^19^F distances to be 9.6 Å for the S1 peak and 14 Å for the S2 and S3 peaks. The PRE-derived distance for the S1 peak is consistent with the distance range between the Ni^2+^ binding site and the ^19^F label calculated from molecular dynamics (MD) simulations of the IFS structure (see **Supplementary Methods**) (mean of 7.7 ± 1.9 Å) (**Fig. 1b**). The PRE-derived distance for the S2 and S3 peaks is significantly larger than for S1 as would be expected for the OFS. It is somewhat smaller than the distance derived from MD simulations of the OFS (mean of 19.4 ± 2.3 Å) (**Fig. 1b**), perhaps because we are beyond the limit of the sensitivity of the measurement (**Fig. 1c**). Collectively, the data allow unambiguous assignment of the S1 peak to the IFS and the S2 and S3 peaks to the OFS.

Detection of two distinct OFS conformations was surprising but not inconsistent with previous findings. Earlier smFRET experiments showed non-homogeneous dynamics of transitions between the OFS and the IFS. The pertinent observation was that the dwell time distribution of the transporter in the OFS was anomalously broad, suggesting that multiple conformations of the OFS existed with different dynamic properties ^17^. Furthermore, crystal structures elucidated an OFS conformation of the transporter but also captured a so-called outward-facing intermediate state (iOFS) of a GltPh mutant, in which the transport domain shifted inward somewhat ^47^. Intrigued by the NMR results, we used Cryo-EM to visualize conformations sampled by GltPh. Toward this end, we imaged wild type GltPh reconstituted into nanodiscs in the presence of saturating concentrations of L-asp and Na^+^ ions (**Supplementary Table S2**). To assess whether there were conformational differences between individual transporter protomers, we performed 3D classification and preliminary refinement using 342,356 particles and applying C3 symmetry (**Supplementary Fig. S3**) followed by symmetry expansion. The focused 3D classification without alignments revealed two structural classes (Class 1 and 2) that were then refined and postprocessed to the final resolution of 3.1 and 3.6 Å, respectively. Notably, we observed no protomers in the IFS, which is consistent with membrane environment favoring the OFS of the transporter compared to detergent micelles ^41^. Class 1 was structurally indistinguishable from the OFS of the transporter (**Figs. 3e and Supplementary S4**) ^45^, while Class 2 closely resembled the iOFS (**Fig. 3e and Supplementary S4**) ^47^. Interestingly, we found a well-structured lipid molecule inserted into a crevice between the transport and scaffold domains in the OFS, but in the iOFS, it was either absent or disordered. To assign these OFS and iOFS states to our two OFS ^19^F NMR signals, we considered the effects of the K290A mutation on the resonance intensities. The mutation, which eliminates a salt bridge in the OFS and should therefore destabilize it, increased the intensities of peaks S1 and S3 relative to peak S2 in the spectrum of K290A/dHis/M385C-TET GltPh protein (**Fig. 2e**). We therefore posit that peak S2 in the ^19^F NMR spectra of the transporter corresponds to the OFS, while peak S3 corresponds to the iOFS. The distance between the ^19^F probe and the bound Ni^2+^ ion is decreased by ∼3 Å in the iOFS structure compared to the OFS, but remains consistent with our PRE measurements.

### Conformational exchange rates

We next performed PRE measurements on the K290A and R276S/M395R (RSMR) mutants, which exchange more rapidly between the IFS and OFS than the wild type GltPh ^17,41^. We observed weaker PREs for the S1 peak of K290A and the RSMR than for the wild type GltPh, and, conversely, stronger PREs for the S2 and S3 peaks (**Fig. 4a, b, Fig. 5** and **Supplementary Table S1**). In contrast, when we repeated the PRE measurements for the K290A mutant bound to the blocker TMA, which prevents transport, we observed PREs that were similar to those measured for the wild type transporter (**Fig. 4b** and **Supplementary Table S1)**, indicating that the conformations of the different states are not significantly perturbed by these mutations. Therefore, the altered PRE effects observed in the more dynamic mutants are likely due to chemical exchange between the peaks.

**Figure 4:**
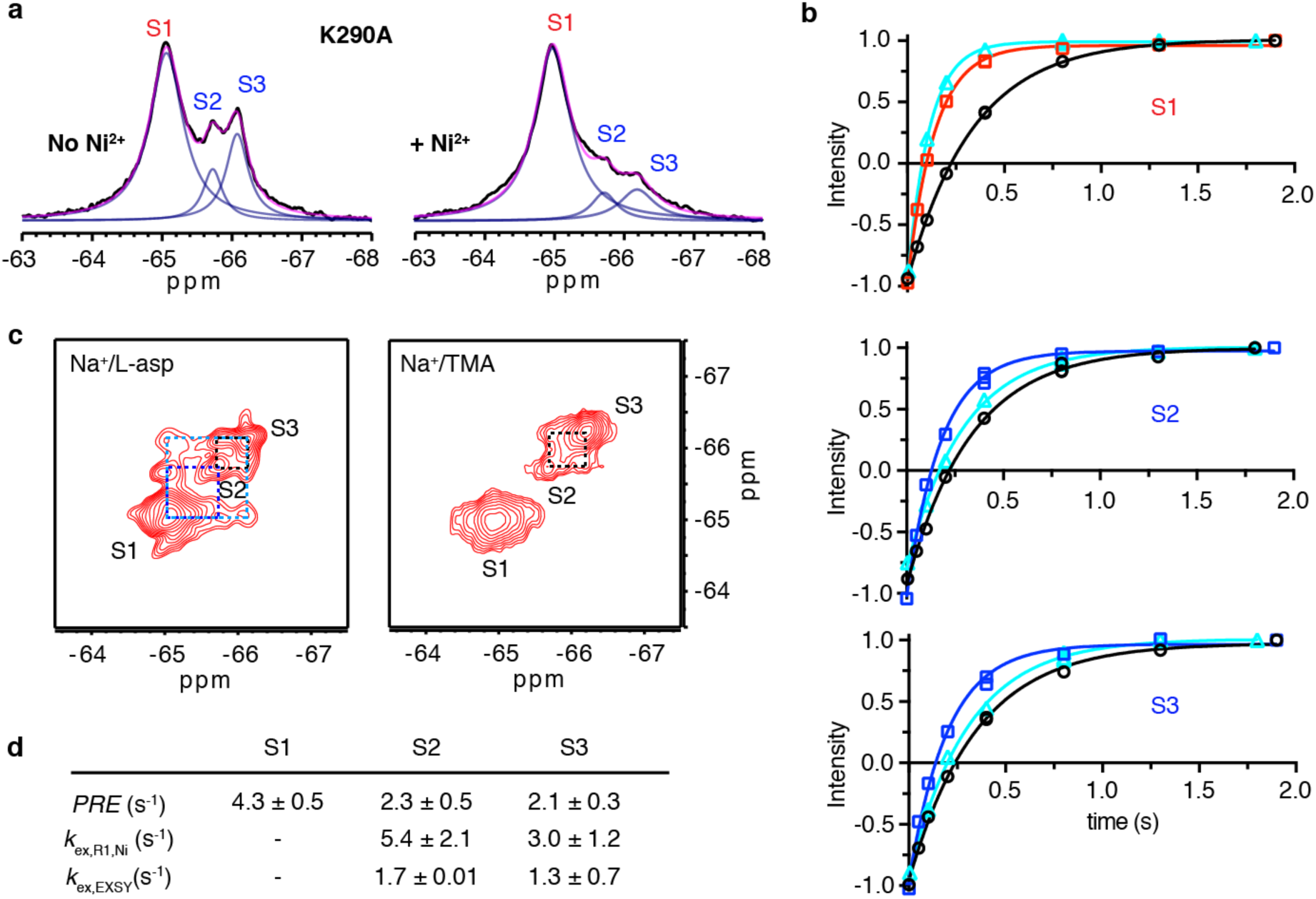
Paramagnetic *R*_1_ relaxation and conformational exchange of K290A mutant. **a**, 1D ^19^F spectra of K290A/dHis/M385C-TET GltPh in the absence (left) and presence (right) of Ni^2+^ ions; **b**, *R*_*1*_ relaxation of S1 (top), S2 (middle) and S3 (bottom) in the absence of Ni^2+^ ions (black), or in the presence of Ni^2+^ and bound to L-asp (red for S1, blue for S2 and S3) or TMA (cyan). Solid lines represent mono-exponential fits with fitted *R*_*1*_ values shown in **Supplementary Table S1. c**, ^19^F-^19^F EXSY spectra of K290A/dHis/M385C-TET GltPh bound to L-asp (left) or TMA (right). Mixing time was set to 0.4 s. Dashed lines indicate cross peaks for S1 and S2 (blue), S1 and S3 (green) and S2 and S3 (black). **d**, Summary of PREs for all peaks and exchange rates between S1 and S2 and between S1 and S3 derived from *R*_*1,Ni*_ or from ^19^F-^19^F EXSY in the presence of L-asp.

**Figure 5:**
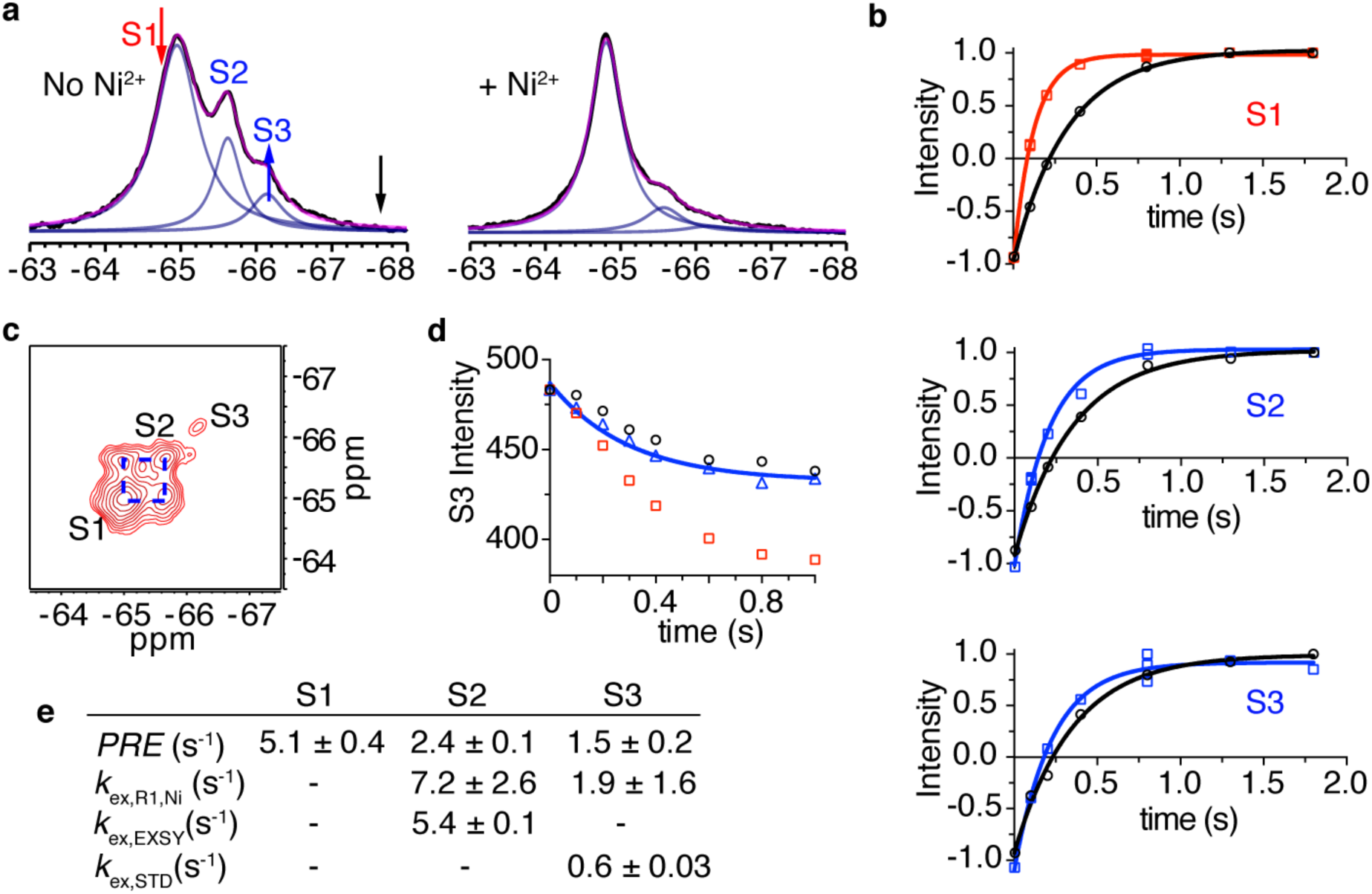
Paramagnetic *R*_1_ relaxation and conformational exchange of RSMR mutant. **a**, 1D ^19^F NMR spectra in the absence (left) and presence (right) of Ni^2+^ ions. The red, black and blue arrows indicate the saturation pulse, control pulse and observed peak, respectively, in the STD experiment in panel **d. b**, *R*_*1*_ relaxation rate of S1 (top), S2 (middle) and S3 (bottom) peaks in the absence (black) and presence (red for S1 and blue for S2 and S3) of Ni^2+^ ions. Solid lines represent mono-exponential fits with fitted *R*_*1*_ values shown in **Supplementary Table S1**. All measurements are in the presence of 200 mM NaCl and 10 µM L-asp. **c**, ^19^F-^19^F EXSY spectrum of RSMR /dHis/M385C-TET GltPh in the presence of 200 mM NaCl and 10 µM L-asp recorded with mixing time of 0.4 s. **d**, Decay of the S3 peak upon saturating the S1 peak (red arrow in **a**) in the STD experiment (red squares). To account for the off-resonance saturation effect, a control experiment (black circles) was performed at an equidistant frequency to S3 peak (black arrow in **a**). The effective decay curve (blue triangle) is fit to Supplementary equation S25, with results given in **Supplementary Table 1. e**, Summary of the PRE values and the exchange rates compared to those determined from EXSY and STD experiments.

Our *R*_1_ measurements in the presence of Ni^2+^ ions can be used to estimate the rate of this chemical exchange. Considering a two-state model where states A and B have ^19^F resonances with strong and weak PREs, respectively,

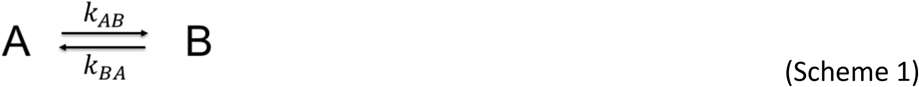

the relaxation processes of the A and B resonances are expected to be bi-exponential because the spins exchange between the two environments (**Supplementary equations S10, S11**). The time evolution of the magnetization of peak B, depends on the intrinsic relaxation rates of the spin in states A and B (*R*\* _*1,A*_ and *R*\* _*1,B*_, respectively) on the populations of the states (*f*_*A*_ and *f*_*B*_) and on the microscopic rate constants (*k*_*AB*_ and *k*_*BA*_). We measured the intrinsic rates *R*\* _*1,A*_ and *R*\* _*1,B*_ separately in the presence of the blocker TMA and the populations of spins in states A and B were obtained by integrating peaks in 1D ^19^F-NMR spectra. Therefore, the relaxation curve of state B can be fitted to the following equation with only two unknown parameters:

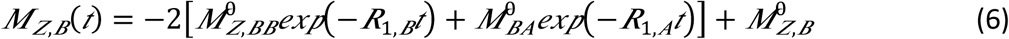

Where *R*_1,*A*_ and *R*_1,*B*_ and 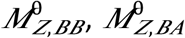 and 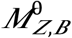 are defined in the **Supplementary Information**.

For the wild type dHis/M385C-TET GltPh, the *R*_*1*_ *PREs* measured for each of the three peaks under paramagnetic conditions were similar in the presence of saturating L-asp or of the blocker TMA. Therefore, as expected, the L-asp bound wild type transporter underwent only slow exchange between the OFS and the IFS with negligible exchange contribution to the *R*_*1*_ *PREs*. To confirm this conclusion, we employed ^19^F-^19^F EXSY to estimate the exchange rates. For the wild type dHis/M385C-TET GltPh, we observed only very weak cross peaks and only between the S1 and S2 peaks at 25 °C (**Supplementary Fig. S5**). Therefore, the OFS and IFS exchange slowly, with the upper limit of the rate estimated at ∼ 0.3 s^−1^, consistent with earlier measurements ^17,41,42^.

In the more dynamic K290A mutant, we considered pairwise equilibria between peaks S1 and S2 and between peaks S1 and S3. S1 (IFS) corresponds to the high-*PRE* state A, while S2 (OFS) and S3 (iOFS) correspond to the low-*PRE* state B in Scheme 1. We then fitted *R*_*1*_ relaxation curves for peaks S2 and S3 using Equation 6 and obtained exchange rates, defined as *k*_*AB*_+*k*_*BA*_, of 5.4 ± 2.1 and 3.0 ± 1.2 s^−1^, respectively (**Fig. 4d, Supplementary Table S1**). For comparison, ^19^F-^19^F EXSY experiments without Ni^2+^ ions resolved cross peaks between all three resonances (**Fig. 4c**). Notably, when EXSY spectra were obtained in the presence of the blocker TMA, cross peaks between S1 and S2 peaks and between S1 and S3 peaks were abolished, while the cross peaks between S2 and S3 peaks were still observed. These results are consistent with the assignment of the S2 and S3 peaks to the OFS and iOFS, and of the S1 peak to the IFS. The exchange rates obtained from EXSY experiments were two to three times smaller than those from the paramagnetic *R*_1_ relaxation measurements (**Fig. 4d**). The differences seem to be due mostly to the faster transition rates from the OFS and iOFS (peaks S2 and S3) to the IFS (peak S1) when bound to Ni^2+^ ions (**Supplementary Table S1**).

For the RSMR mutant, the exchange rates determined from paramagnetic *R*_*1*_ measurements were 7.2 ± 2.6 s^−1^ between peaks S1 and S2 peaks and 2 ± 1.6 s^−1^ between peaks S1 and S3 (**Fig. 5b** and **e**). Because of the low population of the S3 peak in the presence of Ni^2+^ ions, *R*_1_ relaxation measurements were of lower precision, but still suggested that the chemical exchange occurred. EXSY spectra also showed large cross peaks between peaks S1 and S2 with an exchange rate of 5.4 ± 0.1 s^−1^ (**Fig. 5c**). Peak S3 was poorly populated and we could not detect cross peaks. Instead, we used the saturation transfer difference (STD) measurement to estimate the exchange rate between peaks S1 and S3 at ∼0.6 s^−1^ (**Fig. 5d**).

When chemical exchange rates are significantly faster than the intrinsic relaxation rate of the high-*PRE* state *R*_1,A_^*^, the measured *R*_1_ rate becomes the same for states A and B, and the observed *R*_1_ is uninformative about the exchange rate (**Fig. 6a** and **Supplementary Equation S24**). Therefore, the upper limit of the measurable chemical exchange rate depends on the intrinsic *R*_*1,A*_^*\**^ rate of the high-*PRE* state and, to a lesser extent, on the populations of the states, *f*_*A*_ and *f*_*B*_. In our experiments on dHis/M385C-TET GltPh in complex with Ni^2+^ ions, the *R*_*1,A*_^*\**^ rate of the high-*PRE* state is ∼9 s^−1^, and faster chemical exchange rates would not be measurable (**Fig. 6a**). However, *R*_*1,A*_^*\**^ relaxation rates increase steeply as the distance between the paramagnetic ion and the ^19^F nuclei decreases. For example, if the distance in the high-PRE state were 6 Å, *R*_1,*A*_^*^ one would expect the *R*_1_^*^ for the state to approach 100 s^−1^, allowing measurements of correspondingly faster exchange rates (**Fig. 6a**). To test this, we moved the labeling site from M385C to A381C, where the distance between ^19^F nucleus and the chelated Ni^2+^ ion in the IFS is expected to be ∼6 Å based on the crystal structure ^48^. We recorded ^19^F spectra of dHis/A381C-TET GltPh and observed three peaks similar to those observed for dHis/M385C-TET GltPh but with greater overlap (**Fig. 6b**). There was also an additional peak further downfield, which we termed S0. In the presence of 3 molar equivalents of Ni^2+^ ions, the populations of peaks S0 and S1 increased relative to peaks S2 and S3, consistent with our results for dHis/M385C-TET GltPh and suggesting that S0 represents an IFS variant. PRE measurements in the presence of the blocker TMA revealed dramatically faster *R*_1,*A*_^*\**^ rates of ∼125 and 98 s^−1^ for peaks S0 and S1, which correspond to distances between Ni^2+^ ion and ^19^F nucleus of 5.8 and 6.1 Å, respectively, similar to the expected distances for the IFS. Thus, strategic positioning of the metal-chelating dHis motif relative to the ^19^F probe can be used to adjust the intrinsic high-PRE *R*_1,*A*_^*\**^ relaxation rate to tune the range of the measurable conformational exchange rates.

**Figure 6:**
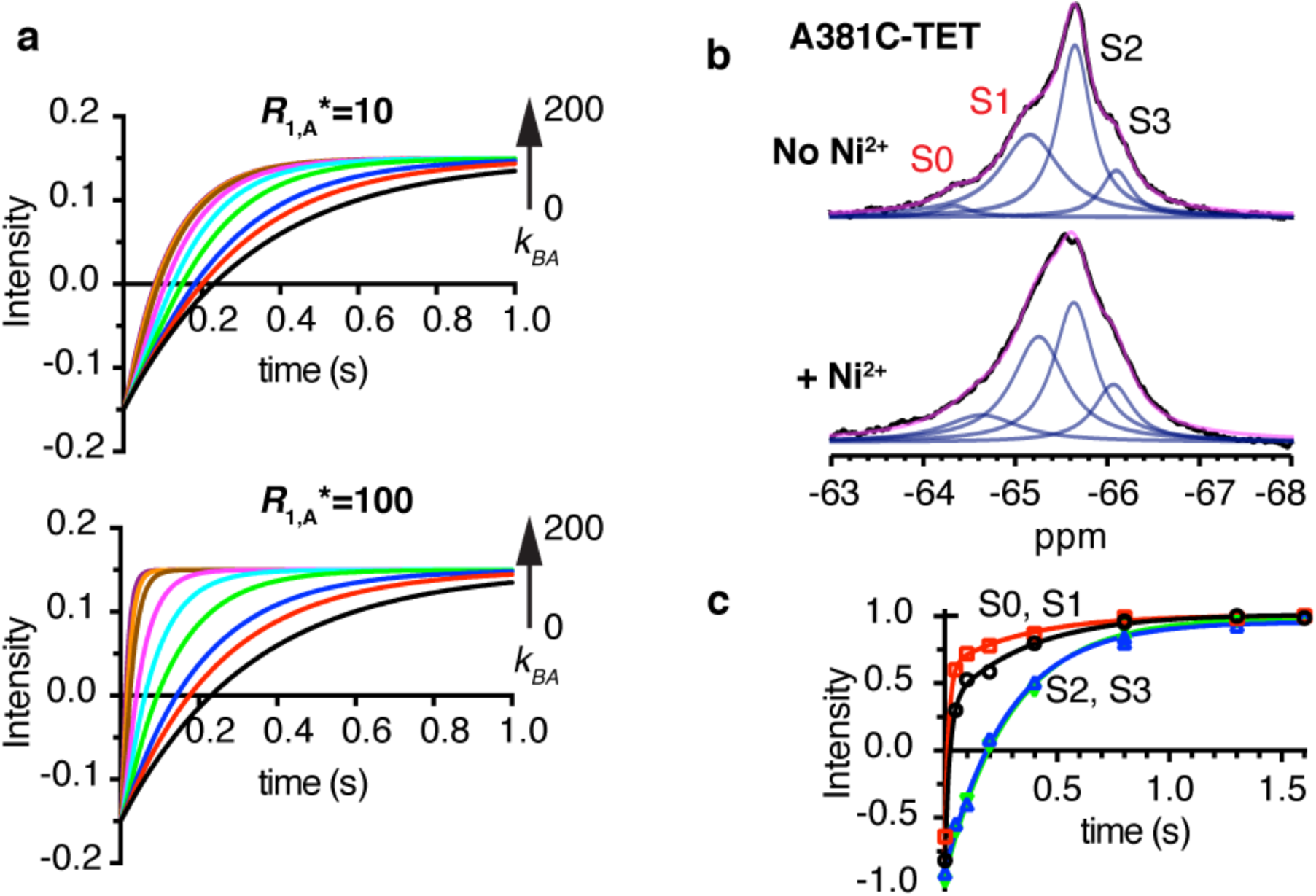
Intrinsic relaxation rate *R*_1,A_^*^ determines the range of accessible exchange rates. **a**, Simulated *R*_1_ relaxation curves of spin B (*R*_1,B_^*^ = 3.0 s^−1^) in exchange with A with *R*_1,A_^*^ = 10 s^−1^ (top) or 100 s^−1^ (bottom). The fraction of spins in state B was set to 0.15 and the rate of transition from state B to state A, *k*_*BA*_, varied from 0 to 200 s^−1^ (black 0, red 1, blue 2, green 5, cyan 10, magenta 20, brown 50, orange 100, purple 200). **b**, 1D ^19^F NMR spectra of dHis/A381C-TET GltPh in the absence (top) and presence (bottom) of 3 molar equivalents of Ni^2+^ ions. **c**, Paramagnetic *R*_*1*_ relaxation curve of the S0 (black circles), S1 (red squares), S2 (blues triangles) and S3 (green reverse triangles) peaks of dHis/A381C-TET GltPh in the presence of Ni^2+^ ions. All measurements were performed in the presence of 200 mM NaCl and 10 µM L-asp. Solid lines represent bi-exponential fits for the S0 and S1 peaks and mono-exponential fits for the S2 and S3 peaks. The fitted parameters for S0 are: *k*_*fast*_ = 124.7 ± 32.0 s^−1^, *k*_*slow*_ = 2.2 ± 1.7 s^−1^; S1: *k*_fast_ = 97.5 ± 5.4 s^−1^, *k*_slow_ = 2.8 ± 0.5 s^−1^; S2: *k* = 5.4 ± 0.8 s^−1^; S3: *k* = 4.5 ± 0.5 s^−1^. The bi-exponential nature of *R*_1_ relaxations of peaks S0 and S1 may reflect the presence of small but significant chemical exchange with the low-PRE peaks.

## Discussion

1D ^19^F NMR of dHis/M385C-TET GltPh bound to Na^+^ ions and L-asp showed three resonances, S1, S2 and S3, which we interpret to correspond to three distinct conformations of the transporter that do not rapidly exchange with each other on the ^19^F NMR time scale. Measurements of Ni^2+^-mediated *PRE*s allowed us to unambiguously assign peak S1 as an IFS and peaks S2 and S3 as two OFS conformations. The effects of mutations and ligands on the observed populations are consistent with these assignments. Our spectra alone do not provide structural details of the observed states, but they can guide high-resolution structural studies employing other techniques. Thus, prompted by our NMR results, we examined GltPh structure by Cryo-EM and observed the presence of two outward-facing states: the OFS and iOFS, to which we assign peaks S2 and S3, respectively. While the populations of the states might not be accurately estimated from Cryo-EM imaging, it is notable that Class 1 particles corresponding to the OFS were more abundant than Class 2 particles, corresponding to iOFS. Consistently, higher populations of the peak S2 (OFS) compared to the peak S3 (iOFS) are observed in ^19^F NMR spectra. NMR can also identify mutations or ligands that favor specific states to facilitate their characterization via other approaches. For example, while both TBOA and TMA are nontransportable blockers ^40,45^, our experiments show that TMA does not have a conformational preference, while TBOA shifts the distributions strongly toward OFS conformations. Similarly, Na^+^ ions also shift the wild type and all mutants examined toward the outward-facing states (**Supplementary Fig. S6**).

Ni^2+^-mediated PRE has the potential to serve as a “molecular ruler”, yielding distances between the metal ion and the ^19^F nucleus. Other methods, such as smFRET and double electron-electron resonance (DEER), can also measure distances between fluorophore or spin labels under favorable circumstances. Unlike these methods, which are most useful in the 20 to 80 Å distance range, *R*_1_ *PRE*-based measurements are most applicable to shorter distances, under 20 Å, making them complementary to other approaches. In many membrane proteins, the shorter distance range is advantageous because the proteins are relatively small, making it difficult to identify suitable labeling sites separated by the requisite longer distances. Distances estimated from our *R*_1_ *PRE* measurements were in reasonable agreement with the values extracted from molecular dynamic simulations of the corresponding states based on existing crystal structures. Nevertheless, there are several factors that may introduce errors into the distance estimates. First, the distances are affected by the rotamers sampled by the TET probe. Similar issues are present in DEER measurements, where frozen samples are analyzed and all sampled rotamers of the spin probes contribute to the observed distance distributions, and in FRET measurements, where preferred orientation of the fluorophores can result in significant changes to the measured distances. In NMR measurements, rotamers are sampled on fast time scales compared to the scale of the experimental measurement and an averaged position of the probe is obtained. Absolute distances calculated from Equation 1 are also affected by the uncertainty in the order parameter, *S*^*2*^, and the internal motion time, τ_*i*_, of the TET probe. This uncertainty stems in part from the fact that these have not been independently determined for the TET side chain, but rather approximated using values typical for methionine.

Comparing the magnitudes of the Ni^2+^-mediated PREs in the wild type dHis/M385C-TET GltPh to those in more dynamic mutants, we observed that in the latter, the PREs for peaks S2 and S3 were larger in the presence of the substrate L-asp, but not when the transporters were bound to the blocker TMA. These increased PREs for the OFS resonances of the dynamic mutants are attributed to the chemical exchange with the IFS, which shows a stronger PRE. From the paramagnetic *R*_1_ measurements, we obtained the pairwise chemical exchange rates between the resonances with high and low PREs, which agree well with those estimated from EXSY and STD. The advantages of paramagnetic *R*_*1*_ measurements are rapidity compared to methods relying on 2D spectra and greater tolerance to spectral overlap. This is critical for studying large molecular assemblies in which NMR resonances tend to be broad and poorly resolved. Thus, PRE-based detection of conformational dynamics provides a viable method to screen for the modulators of membrane transporters, channels and receptors.

Our analysis shows that both OFS and iOFS conformations of the K290A and RSMR mutants of GltPh exchange with the IFS. However, it is not possible to unambiguously establish the order of the events. Thus, the OFS conformation may either transition first into the iOFS and from there into the IFS, or alternately transition directly into the IFS with rare visits of the iOFS state. Interestingly, the exchange rates between the S1 (IFS) and S2 (OFS) peaks are higher than between the S1 and S3 (iOFS) peaks for both mutants. This may suggest that iOFS is not an obligatory intermediate between the OFS and the IFS states. Notably, data interpretation for GltPh might be further complicated by the kinetic complexity revealed by single molecule FRET recordings for this transporter ^41^ but may be masked by ensemble averaging in ^19^F NMR experiments.

In summary, our method provides an avenue to study both equilibrium distributions and dynamics of multiple conformational states of membrane proteins in solution. It should be broadly applicable to the study of membrane proteins undergoing conformational transitions on sub-second to second time scales.

## Acknowledgements

The authors thank Dr. X. Yao for helpful discussions, Dr. M. Goger and Dr. S. Bhattacharya for help with setting up NMR, and Dr. M. A. Cuendet for help with setting up initial MD simulations. We thank Dr. Chen Xu and Dr. KangKang Song at the UMass cryo-EM facility for help with electron microscopy data collection. This work was supported by NIH grants R37NS085318 and R01NS064357 (OB), R37AG019391 (DE) and S10OD016320 (CB). OB, DE and CB are members of the New York Structural Biology Center (NYSBC) which is supported in part by NIH Grant P41 GM118302 (CoMD/NMR: Center on Macromolecular Dynamics by NMR Spectroscopy), ORIP/NIH facility improvement grant CO6RR015495 and NIH grant S10OD018509. The coordinates of the structures and the density maps have been submitted to Protein Data Bank under the accession codes of 6UWF and 6UWL.

## Author contributions

Y.H., D.E. and O.B. designed the experiments; Y.H. and G.L. performed the NMR experiments; Y.H. and O.B analyzed the data; C.B. assisted with NMR experimental design, data collection and analysis; X.W. performed Cryo-EM imaging, analyzed data and refined molecular models; G.H. performed transport activity assays; A.M.R. performed molecular dynamics simulations; Y.H., X.W., A.M.R., O.B., D.E., H.W. and C.B. wrote the manuscript.

## Supplementary Information

### Theory

The magnitude of *R*_1_ *PRE* is determined by the dipole-dipole interaction between the spins of the unpaired electrons and of a nucleus as described by the Solomon-Bloembergen (SB) equations ^26,27^:

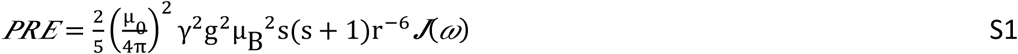

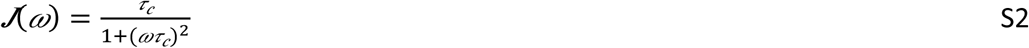

where *μ*_*0*_ is the permeability of vacuum (4π × 10^−7^ m kg s^−2^ A^−2^), *γ* is the nuclear gyromagnetic ratio (*γ* _*F*_=25.166 × 10^−7^, *γ*_*H*_ =26.752 × 10^−7^), g is the electron Land g-factor (−2.0023193), *μ*_*B*_ is the magnetic moment of the free electron (−9.284764 × 10^−24^ J/T), s is the electron spin quantum number (see **Fig. 1d** for examples), *r* is the distance between the electron and the nucleus, and *ω* is the nuclear Larmor frequency (4π × 470 × 10^6^ rad/s for ^19^F and 4π × 500 × 10^6^ rad/s for ^1^H in the instrument we use).

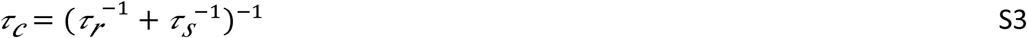

where τ_*r*_ is the isotropic protein rotation correlation time and τ_*s*_ is electron relaxation time. For a protein with hydrodynamic radius *R*, τ_*r*_ can be estimated using Stoke’s law:

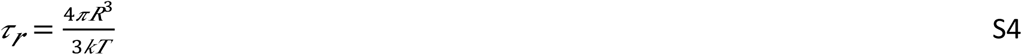

where *k* is the Boltzmann constant (1.3806 × 10^−23^ m^2^ kg s^−2^ K^−1^) and *T* is the absolute temperature. For a ∼300 kDa protein/detergent particle with hydrodynamic radius of 59 Å we estimate τ_*r*_ to be 213 ns. Electron relaxation times, τ_*s*_, for various paramagnetic centers are shown in **Figure 1d**.

To take into an account the local motion, we expand the SB equation using the model-free approach as previously described ^28,29,31^:

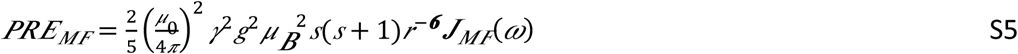

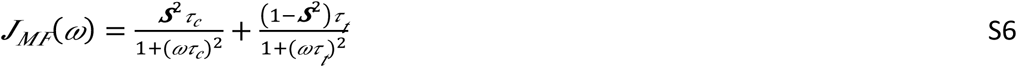

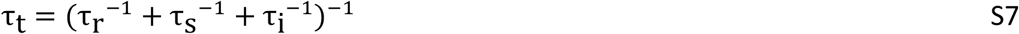

τ_*i*_ is the internal correlation time of the ^19^F label and *S*^*2*^ is the order parameter. For TET label, we use τ_*i*_ of 20 ps and *S*^*2*^ of 0.1 measured previously for the methionine side chain ^36,37^ as an approximation.

To estimate the effect of the chemical exchange on the paramagnetic *R*_1_ relaxation, we consider a spin in chemical exchange between a state A with strong PRE and a state B with weak PRE:

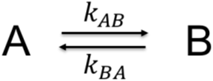

The time evolution of the longitudinal magnetizations for the two states, *M*_*Z,A*_ and *M*_*Z,B*_ is described by the modified McConnell equations ^53^, which are provided below for clarity:

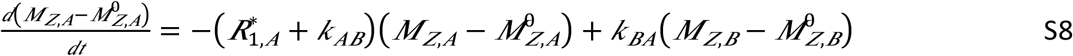

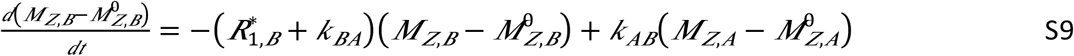

where *M*^*0*^_*Z,A*_ and *M*^*0*^_*Z,B*_ are the magnetizations at time 0 for states A and B, respectively, and *R*^*\**^_*1,A*_ and *R*^*\**^_*1,B*_ are the intrinsic relaxation rates of the spins in these states. For an inversion recovery experiment, under the initial condition 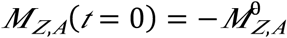 and 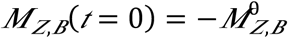 the solutions are:

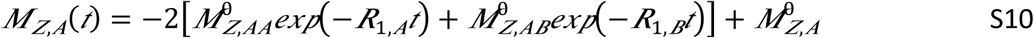

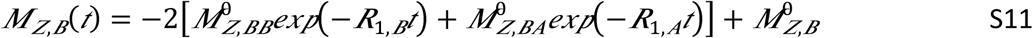

Note that

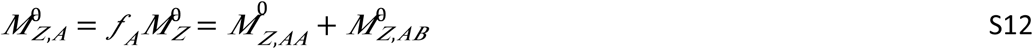

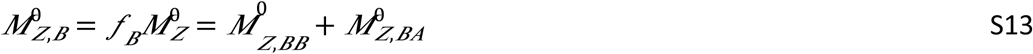

where

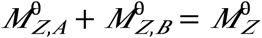, is the total initial magnetization

and the coefficients 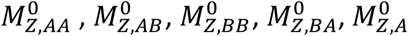 and 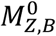 are:

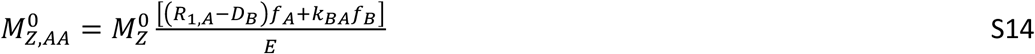

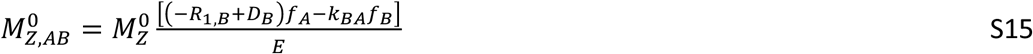

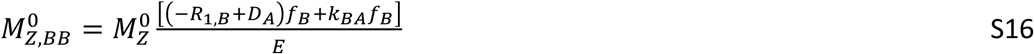

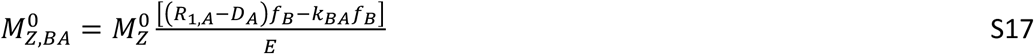

*f*_*A*_ and *f*_*B*_ are equilibrium fractions of states A and B. *R*_*1,A*_ and *R*_*1,B*_ are relaxation rates of the fast and slow phase of the longitudinal relaxation curve in the presence of the chemical exchange, respectively, ^53,54^. They depend on the equilibrium fractions of states A and B, on the transition rate *k*_*BA*_, and on the intrinsic relaxation rates of the spin, *R*^*^_*1,A*_ and *R*^*\**^_*1,B*_:

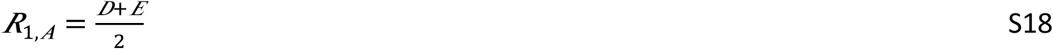

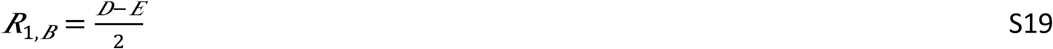

where

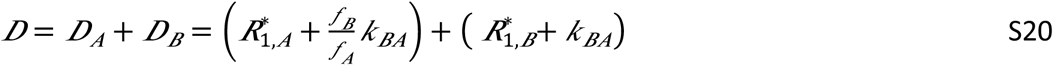

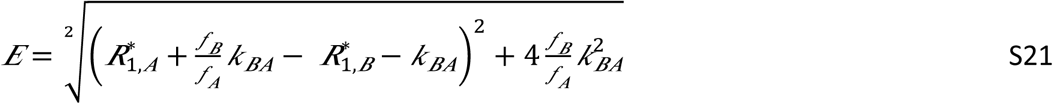

The intrinsic relaxation rates *R*\* _*1,A*_and *R*\* _*1,B*_, are measured separately in the presence of the blocker, and the values of *f*_A_ and *f*_B_ are obtained by integrating deconvoluted peaks in 1D ^19^F-NMR spectra. Therefore, fitting *R*_*1,B*_ relaxation curves for spin B to equations S11 requires optimization of only two parameters: *k*_BA_ and 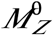.

Notably, if exchange is very slow, (i.e., 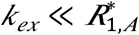), 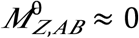 and 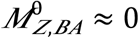 the *R*_1_ relaxation becomes mono-exponential:

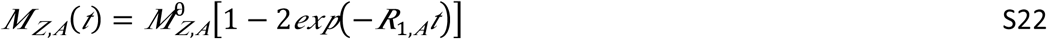

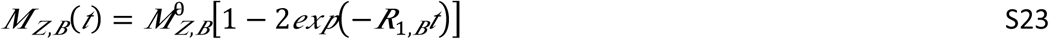

i.e. spins in states A and B relax with rate *R*_1,*A*_ and *R*_1,*B*_, respectively.

In the case of the fast exchange, (i.e., 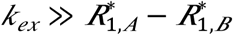) ^55^, 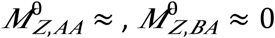, and states A and B relax with the same rate *R* _1,*B*_, and

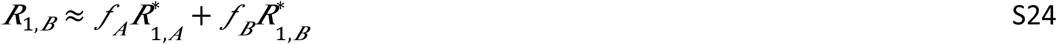

## Supplementary Methods

### Protein expression, labeling and purification

The fully functional cysteine-free GltPh C321A with seven additional histidines, which significantly improve expression yield ^56^, is referred to the wild type throughout the paper for brevity. M385C and other mutations were introduced using QuikChange kit (*Qiagen*). The protein constructs with carboxyl-terminal thrombin cleavage site and (His)_8_-tag were purified as described previously ^56^. Briefly, the plasmids were transformed into *E. coli* DH10-B cells (*Invitrogen*). Cells were grown in LB media supplemented with 0.2 mg/l of Ampicillin (*Goldbio*) at 37 °C until OD_600_ of 1.0. Protein expression was induced by adding 0.2 % arabinose (*Goldbio*) for 16 hr at 24 °C. The cells were harvested by centrifugation and re-suspended in 20 mM Hepes, pH7.4, 200 mM NaCl, 1 mM L-asp, 1 mM EDTA. The suspended cells were broken by Emulsiflex C3 high pressure homogenizer (*Avestin Inc*.) in the presence of 0.5 mg/ml lysozyme (*Goldbio*) and 1 mM Phenylmethanesulfonyl fluoride (PMSF, *MP Biomedicals*). After centrifugation for 15 min at 5000 g at 4 °C to remove the debris, membranes were pelleted by centrifugation at 125000 g for 60 min. The membranes were homogenized in 20 mM Hepes, pH 7.4, 200 mM NaCl, 1 mM L-asp, 10 mM EDTA, and 10 % sucrose. The suspension was centrifuged at 125000 g for 60 min. The crude membranes were collected and homogenized in Buffer A, containing 20 mM Hepes, pH7.4, 200 mM NaCl, 1 mM L-asp, at 8 ml per gram of membranes. For fluorine labeling ^19^, membranes were solubilized in the presence of 40 mM n-dodecyl-β-D-maltopyranoside (DDM, *Anatrace, Inc*.) and 2 mM 2,2’-dithiodipyridine (DTDP, *Sigma Aldrich*) for 2 hours at 4 °C. The mixture was centrifuged for 60 min at 125000 g, the supernatant was diluted 4 times with Buffer A and incubated with Ni-NTA resin (*Qiagen*) for 1 hour at 4 °C. The resin was washed with 5 volumes of Buffer A with 1 mM DDM. The resin slurry was supplemented with 2 mM trifluoroethanethiol (*Sigma Aldrich*) and incubated with mixing at 4 °C overnight. After washing the resin with 10 volumes of Buffer A containing 1 mM DDM and 25mM imidazole, the labeled protein was eluted in the same buffer containing 250 mM imidazole. The protein samples were concentrated to ca. 10 mg/ml using concentrators with 100 kDa MW cutoff (*Amicon*). Protein concentration was determined by UV absorbance at 280 nm using extinction coefficient of 57400 M^−1^ cm^−1^ and MW of 44.7 kDa. The (His)_8_-tag was cleaved by thrombin (Sigma) using 20 U per 1 mg GltPh in the presence of 5 mM CaCl_2_ at room temperature overnight. The reaction was stopped by addition of 10 mM EDTA and 1 mM PMSF. The protein was further purified by size exclusion chromatography (SEC) on Superdex 200 Increase 10/300 GL column (*GE Healthcare Life Sciences*). GltPh bound to L-Asp condition was prepared using SEC in buffer containing 20 mM Hepes, pH7.4, 100 mM NaCl, 10 µM L-Asp, 1 mM DDM. To prepare substrate-free GltPh, the SEC buffer with 20 mM Hepes, pH 7.4, 50 mM KCl, 1 mM DDM was used. NaCl and ligands in appropriate concentrations were then added into the eluted protein solution. The protein was then concentrated and either used immediately or snap-frozen in liquid nitrogen and then stored at −80 °C. To evaluate the efficiency of TET labeling, labeled and unlabeled protein samples were incubated with 10 molar equivalents of fluorescein-5-malaimide in the presence of 2% SDS for 4h at room temperature. The reaction mixtures were than analyzed by SDS-PAGE followed by fluorescence imaging.

### Transport activity assay

Labelled M385C and dHis M385C GltPh proteins were reconstituted into liposomes and ^3^H L-asp uptake was measured as previously described ^45^. Briefly, liposomes were prepared from a 3:1 (w/w) mixture of *E. coli* total lipid extract and egg yolk phosphotidylcholine (*Avanti Polar Lipids*) in 20 mM Tris/HEPES, pH7.4, and containing 200 mM KCl and 100 mM CholineCl. Liposomes were destabilized by addition of Triton X-100 at detergent to lipid ratio of 0.5:1 (w/w). GltPh proteins were added at final protein to lipid ratio of 1:2000 (w/w) and incubated for 30 min at room temperature. Detergents were removed by repeated incubations with Bio-Beads™ SM-2 resin (Bio-Rad). The proteoliposomes were subjected to three freeze-thaw cycles and extruded through 400 nm filters before the uptake assay. Uptake reaction was started by diluting the proteoliposomes 100-fold into reaction buffer containing 20 mM HEPES/Tris pH 7.4, 200 mM KCl, 100 mM NaCl, the indicated amounts of L-asp (L-asp concentrations above 1 μM were supplemented with 5 μM cold L-asp, maximally diluting ^3^H-L-asp 10-fold) and 0.5 μM valinomycin. Uptake was performed for 1 min at 35 °C.

### ^19^F NMR spectroscopy

^19^F-NMR experiments were performed using a Bruker Avance IIIHD 500 MHz spectrometer equipped with a TCI ^1^H-^19^F/^13^C/^15^N triple resonance cryogenic probe (*Bruker Instruments*). 50 μM TFA and 10 % D_2_O were included in the sample and used as chemical shift reference (−75.4 ppm) and lock reagent, respectively. 1D ^19^F-NMR spectra were recorded with 4096 points and spectral width (SW) of 40 ppm. Acquisition time was 109 ms under these settings. Carrier frequency was set to −70 ppm. The number of scans was set between 512 and 4096 depending on the sample conditions. The recycle delay was 0.6 s for 1D ^19^F spectra except when otherwise indicated. 90° pulse length was calibrated for each sample and was typically 11.4 μs. All the spectra were recorded without ^1^H decoupling. 1D spectra were processed using MestRaNova 12.0.0 software (*Mestrelab Research*) employing a 20 Hz exponential windows function, zero filling to 16 k points. The spectra were baseline-corrected, the peaks fitted to Lorentzian peak shapes and assessed based on fit residuals.

^19^F longitudinal relaxation rates, were measured by the inversion recovery method. The recycle delay was 1.8 s, which is ∼5 times *T*_1_. Eight different delays between 0.05 ms and 1.8 s were randomly sampled, with one point repeated 3 times. The peak intensities were obtained from spectral deconvolution. The *R*_1_ relaxation rates and standard errors were obtained by fitting data to single exponential function *I* = *I*_0_(1 − 2*exp*(−*R*_1_*t*). The *R*_1_ *PRE*s were obtained by subtracting the relaxation rate measured in the absence of Ni^2+^ ion from the rate measured in the presence of the ion. The errors were calculated by error propagation. All experiments were repeated at least twice.

^19^F-^19^F EXSY experiments were recorded with 4096 points and spectral width of 40 ppm in the direct dimension and with 24 complex points and SW of 5 ppm in the indirect dimension. Recycle delay was 1.2 s. Typically, 512 to 1024 scans per increment were accumulated depending on sample concentrations. The spectra were processed using Topspin 3.5 software (*Brucker Instruments*). The indirect dimension was zero-filled to 128 points and a Gaussian window function was applied with line broadening factor (LB) of −20 Hz and gaussian multiplication factor (GM) of 0.02. In the direct dimension, the GM window function was applied with LB of −20 Hz and GB of 0.01. The processed spectra were analyzed using Sparky software (T. D. Goddard and D. G. Kneller, SPARKY 3, University of California, San Francisco) and the peak intensities were obtained by integration with Lorentzian fit. A spectrum with mixing time was 0.4 s, unless otherwise stated, and a spectrum with 0 s mixing time were recorded. The peak volumes were fitted using EXSYcalc software (*Mestrelab Research*).

^19^F Saturation transfer difference (STD) spectra were recorded using a train of 50 ms 180° Gaussian shaped pulses with total duration of 0, 100, 200, 300, 400, 600 and 1000 ms. For each duration, on and off resonance irradiation pulses (as shown in Figure 5a) were alternately interleaved to generate saturation and control datasets ^57,58^. The corrected intensities of the observed peak A upon saturation duration on peak B were fitted to the following equation to determine *k*_AB_:

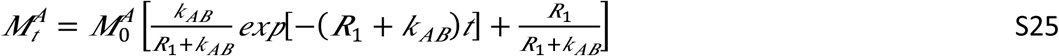

where *R*_1_ is the longitudinal relaxation rate of the observed peak. The backward rate is obtained using the equilibrium populations of A and B, i.e., *k*_BA_ = *k*_AB_^*^*f*_A_/*f*_B_. The fitting was performed in Prism 6 (*GraphPad Software Inc*.)

### Molecular dynamics simulations

The molecular models of the outward-facing GltPh bound to 3 Na^+^ ions and L-asp was constructed from PDB id: 2NWX, and of the inward-facing GltPh bound to 3 Na^+^ ions and L-asp from PDB id: 3KBC. Missing loops and residues were added with the Modeler software package ^59^. The system was immersed in a POPC membrane in a neutralizing solution of NaCl at 0.15 M using the CHARMM-GUI membrane builder web interface ^60^, yielding a simulation box with dimensions: 165 × 175 × 86 Å^3^ and about 250,000 atoms for each system. CHARMM36 parameters ^61^ were used for protein and lipids in all simulations. A standard equilibration protocol developed for various membrane systems ^62^ was applied as follows: the solvent and membrane were equilibrated with harmonic restraint applied to protein and heavy atoms (except those modeled). These restraints were gradually decreased from 20 kcal/mol/Å^2^ to 0.1 kcal/mol/Å^2^ over the course of 5 ns simulation, followed by an additional 13 ns with restraints on the Cα atoms and bound sodium atoms, and an additional 6 ns without any restraints. The last frames of these equilibration phases were used to add the NMR probe and metal ions as follow: M385 was mutated to cysteine modified with NMR probe TET, and residues 215 and 219 were mutated to histidine. The force field parameters for the TET probe, with net zero charge, was adapted from CHARMM generalized force field parameters for difuorobenzylphosphonate and the regular methionine amino acid. To mimic the Ni^2+^ site between H215 and H219, harmonic restraints (10 kcal/mol/Å^2^) were used to restrain Zn^2+^ (with available CHARMM parameters) near these His residues. The systems were then minimized and the final production runs, about 100 ns for each system, were performed using NAMD software version 2.13 ^63^ at 310 K in NPT ensemble with 2 fs time steps. Particle mesh Ewald ^64^ was used for long range electrostatic interactions. Short range electrostatic interactions were calculated using Lennard-Jones potential with truncation distance cutoff at 11 Å with smoothing starting at 8 Å. Pressure was controlled with Langevin barostat at 1 bar with oscillation period of 200 fs and damping time of 100 fs. The X, Y, Z cell dimensions were allowed to fluctuate independently. Temperature was controlled with Langevin thermostat with damping coefficient of 0.1 1/ps. The visualization and quantitative analyses of all simulations are done with VMD ^65^ and MDTraj software ^66^.

### CryoEM structure determination

#### GltPh reconstitution into nanodiscs

Membrane scaffold protein MSP1E3 was expressed and purified from *E. coli* and GltPh was reconstituted into lipid nanodiscs as previously described, with modifications ^67^. Briefly, *E. coli* polar lipid extract and egg phosphatidylcholine in chloroform (*Avanti*) were mixed at 3:1 (w:w) ratio and dried on rotary evaporator and under vacuum overnight. The dried lipid film was resuspended in buffer containing 200 mM NaCl, 1 mM L-asp, 80 mM DDM, 20 mM Hepes/Tris, pH 7.4 by 10 freeze/thaw cycles resulting in 20 mM lipid stock. Purified GltPh protein in DDM was mixed with MSP1E3 and lipid stock at 1:1:50 molar ratio at the final lipid concentration of 5 mM and incubated at 21 °C for 30 min. Biobeads SM2 (Bio-Rad) were added to one third of the reaction volume and the mixture was incubated at 21 °C for 2 hr on a rotator. Biobeads were replaced and incubated at 4 °C overnight. The reconstitution mixture was cleared by centrifugation at 100,000 g and GltPh-containing nanodiscs were purified using a Superose 6 Increase 10/300 GL column (GE Lifesciences) pre-equilibrated with buffer containing 200 mM NaCl, 1 mM L-asp, 20 mM Hepes/Tris, pH7.4.

#### Cryo-EM data collection

To prepare cryo-grids, 3.5 μL of GltPh-containing nanodiscs (6 mg/mL) supplemented with 1.5 mM fluorinated Fos-Choline-8 (*Anatrace*) was applied to a glow-discharged UltrAuFoil R1.2/1.3 300-mesh gold grid (*Quantifoil*) and incubated for 20 s under 100 % humidity at 15 °C. Following incubation, grids were blotted for 2 s and plunge frozen in liquid ethane using a Vitrobot Mark IV (FEI). Cryo-EM imaging data were acquired on a Titan Krios microscope (*Thermo Fisher Scientific*) operated at 300 kV with a K3 Summit direct electron detector (*Gatan*). Automated data collection was carried out in super-resolution counting mode using SerialEM software ^68^ with a magnification of 81,000 x, which corresponds to a calibrated pixel size of 1.06 Å on the specimen and 0.53 Å for super-resolution images. Each movie had a total dose of 50.1615 e^−^/Å^2^ distributed over 30 frames (1.672 e^−^/ Å^2^/frame) with an exposure time of 8.1 s (269 ms/frame) and a defocus range of −1.5 to −2.5 μm.

#### Image processing

The frame stacks were motion corrected using MotionCorr2 ^69^ and contrast transfer function (CTF) estimation was performed using CTFFIND4 ^70^. All processing steps were done using RELION 3.0 unless otherwise indicated ^71^. Dogpicker ^72^ as part of the Appion processing package ^73^ was used for reference-free particle picking. Picked particles were then extracted and subjected to 2D classification to generate 2D class-averages which were used as templates for automated particle picking in Relion. The particles were extracted using a box size of 256 Å with 2x binning and subjected to 2 rounds of 2D classification ignoring CTFs until the first peak. 926,308 particles selected from 2D classification were further classified into 6 classes without applying symmetry using an initial model from a density map previously obtained for a GltPh-containing nanodisc (unpublished) filtered to 40 Å. 342,356 particles from the best class showing a trimeric transporter arrangement were re-extracted, unbinned, and subjected to 3D refinement applying C3 symmetry. After conversion, the refinement was continued with a mask excluding the nanodisc, resulting in a 3.35 Å resolution map. The resolution of the refined map was assessed using Relion postprocessing and gold standard FSC value 0.143 using a mask that excluded the nanodisc. The 3D refinement with C3 symmetry produced a map of GltPh in the OFS with bound substrate closely resembling the PDB model 2NWX. To probe for conformational heterogeneity, we employed the symmetry expansion implemented in Relion ^74^. 1,027,068 protein subunits were rotated to the same position and subjected to a focused 3D classification without alignment with T=40 into 10 classes. The local mask was generated using Chain A of PDB model 2NWX and included only densities from one subunit on the refence map. Nine classes were either low resolution or pictured GltPh in the OFS conformation. One class, which contained 11.7% of symmetry expanded particles, showed a different conformation, closely resembling the iOFS conformation. The best OFS class (94,731 particles) and the iOFS class (120282 particles) were separately subjected to a final focused 3D refinement with C1 using a mask to exclude the nanodisc. The local angular searches in this refinement were conducted only around the expanded set of orientations to prevent contributions from neighbor subunits in the same particle. The resulting maps were postprocessed in Relion using the same mask as in 3D classification after symmetry expansion. The final resolution at gold standard FSC value 0.143 was estimated as 3.1 Å for the OFS map and 3.6 Å for iOFS map. Local resolution variations were estimated using ResMap ^75^.

#### Model building and refinement

For atomic model building, one subunit of 2NWX GltPh crystal structure was docked into the density map of the OFS using UCSF Chimera ^76^. For iOFS, one subunit of 3V8G crystal structure was docked into the density. For each model after the first round of real-space refinement using Phenix ^77^, miss-aligned regions were manually adjusted and missing side chains and residues were manually added in COOT ^78^. Phosphatidylethanolamine is used as a model lipid to be placed into densities which resemble lipid molecules with its acyl chains or ethanolamine heads truncated to fit the visible densities. Models were iteratively refined applying secondary structure restraints and validated using Molprobity ^79^ and EMRinger ^80^. For further cross validation and to check for overfitting, all atoms of each model were randomly displaced by 0.3 Å and each resulting model was refined against the first half-map obtained from processing. FSC between the refined models and the half-maps used during the refinement were calculated and compared to the FSC between the refined models and the other half-maps. In addition, the FSC between the refined model and sum of both half-maps was calculated. The resulting FSC curves were similar showing no evidence of overfitting.

## Supporting Figures and Tables

**Figure S1.**
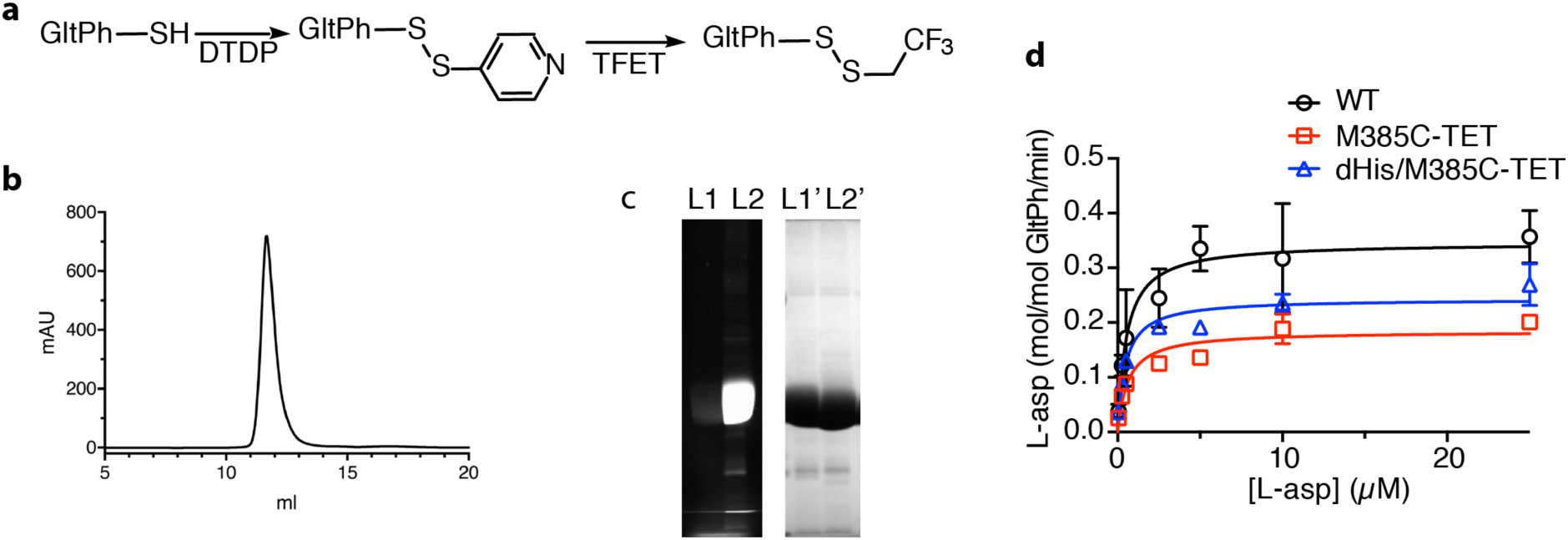
Protein purity, labeling efficiency and ^3^H L-asp uptake. **a**, Scheme for site-specifically introducing ^19^F label into M385C GltPh mutant **b**, Size exclusion chromatography elution profile of M385C-TET GltPh. **c**, SDS-PAGE gel imaged by fluorescence (left) and Coommasie blue staining (right) of M385C GltPh labeled with fluorescein-5-malaimide before (lane 1) and after (lane 2) labeling with TET. Protein samples were incubated with 10-fold excess of fluorescein-5-malamide for 4h prior to analysis. **d**. Michaelis-Menten kinetics of ^3^H L-Asp uptake for wide type (WT) GltPh (cycle), M385C-TET (square), and dHis/M385C-TET GltPh (triangle). DTDP: 2,2’-dithiodipyridine; TFET: trifluoroethanethiol.

**Figure S2.**
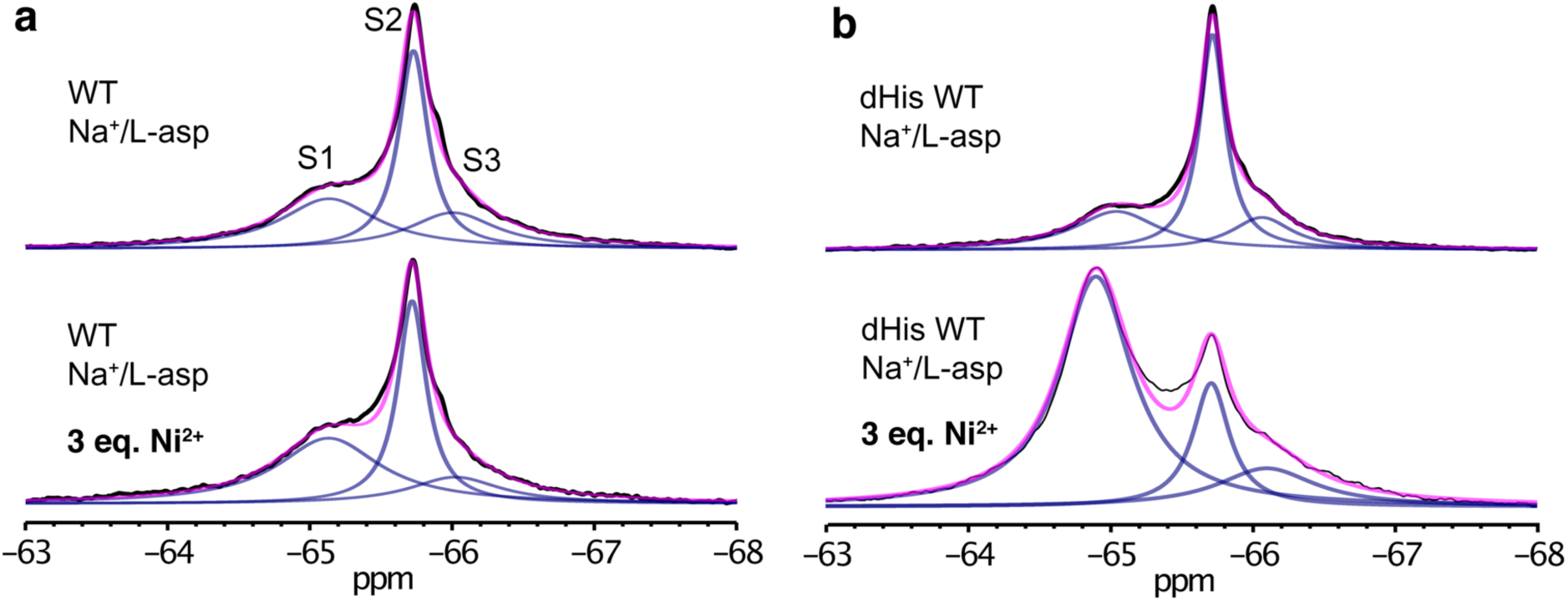
Specific Ni^2+^ binding to dHis/M385C-TET GltPh mutant. 1D ^19^F-NMR spectra of M385C-TET GltPh (**a**) and dHis/M385C-TET GltPh (**b**) without (up) and with (bottom) 3 molar equivalents of Ni^2+^ ions. Spectra were recorded at 293K in the presence of NaCL and L-asp.

**Figure S3.**
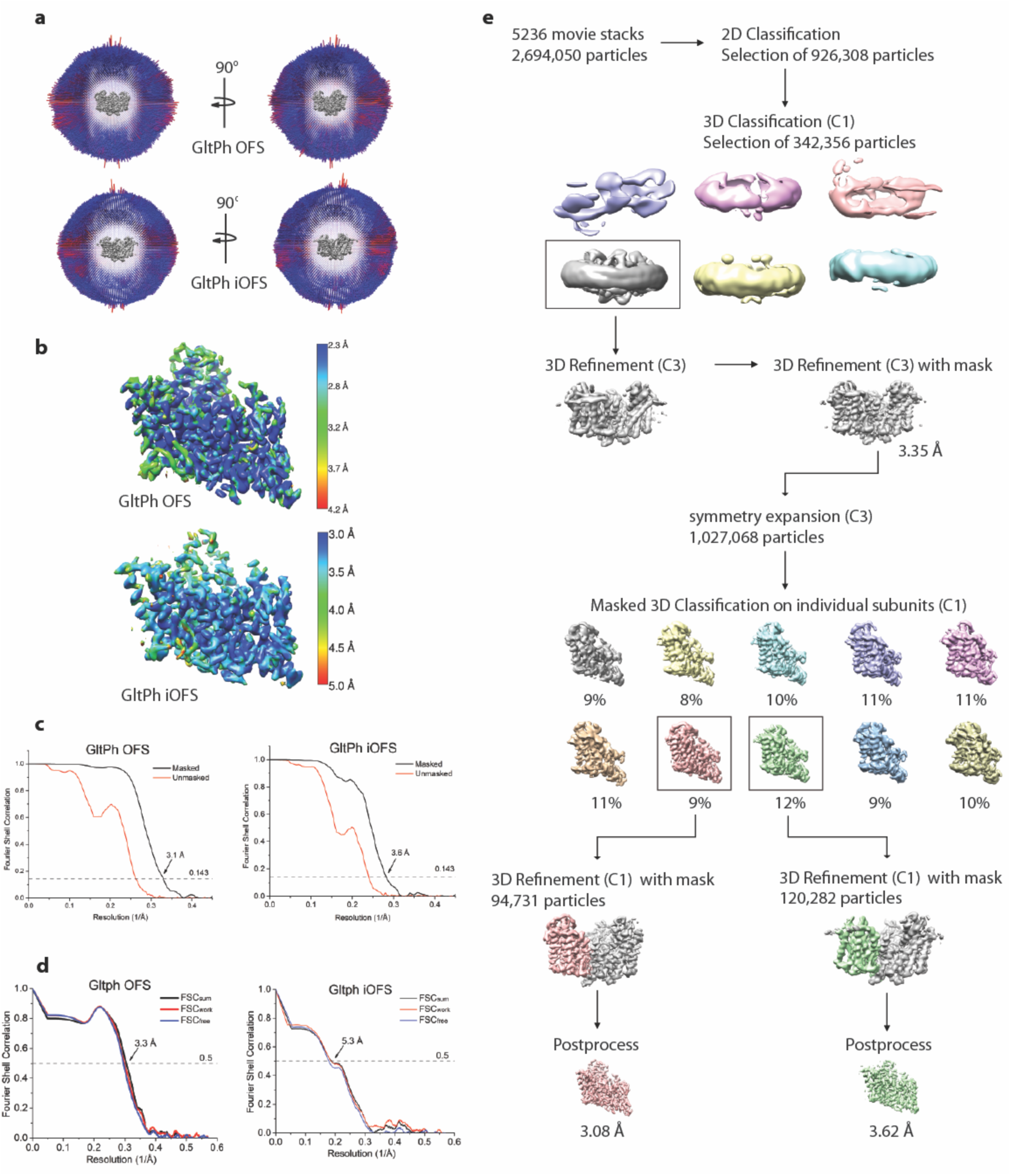
Cryo-EM data processing. **a**, Angular distribution of particles contributing to the final reconstitution. Number of views at each angular orientation is represented by length and color of cylinders where red indicates more views. **b**, Final maps after Relion post-processing colored according to local resolution estimation using ResMap. **c**, Fourier shell correlation (FSC) curves indicating the resolution at the 0.143 threshold of final masked (black) and unmasked (orange) maps of GltPh OFS (left) and iOFS (right). **d**, FSC curves from cross validation of refined GltPh OFS (left) and iOFS (right) models compared to the masked half-map 1 (Orange traces: FSC_work_, used during validation refinement), masked half map 2 (Blue traces: FSCf_ree_, not used during validation refinement), and the masked summed map (Black traces: FSC_sum_). **e**, Data processing flow chart for GltPh reconstituted into nanodisc in the presence of NaCl and L-asp.

**Figure S4.**
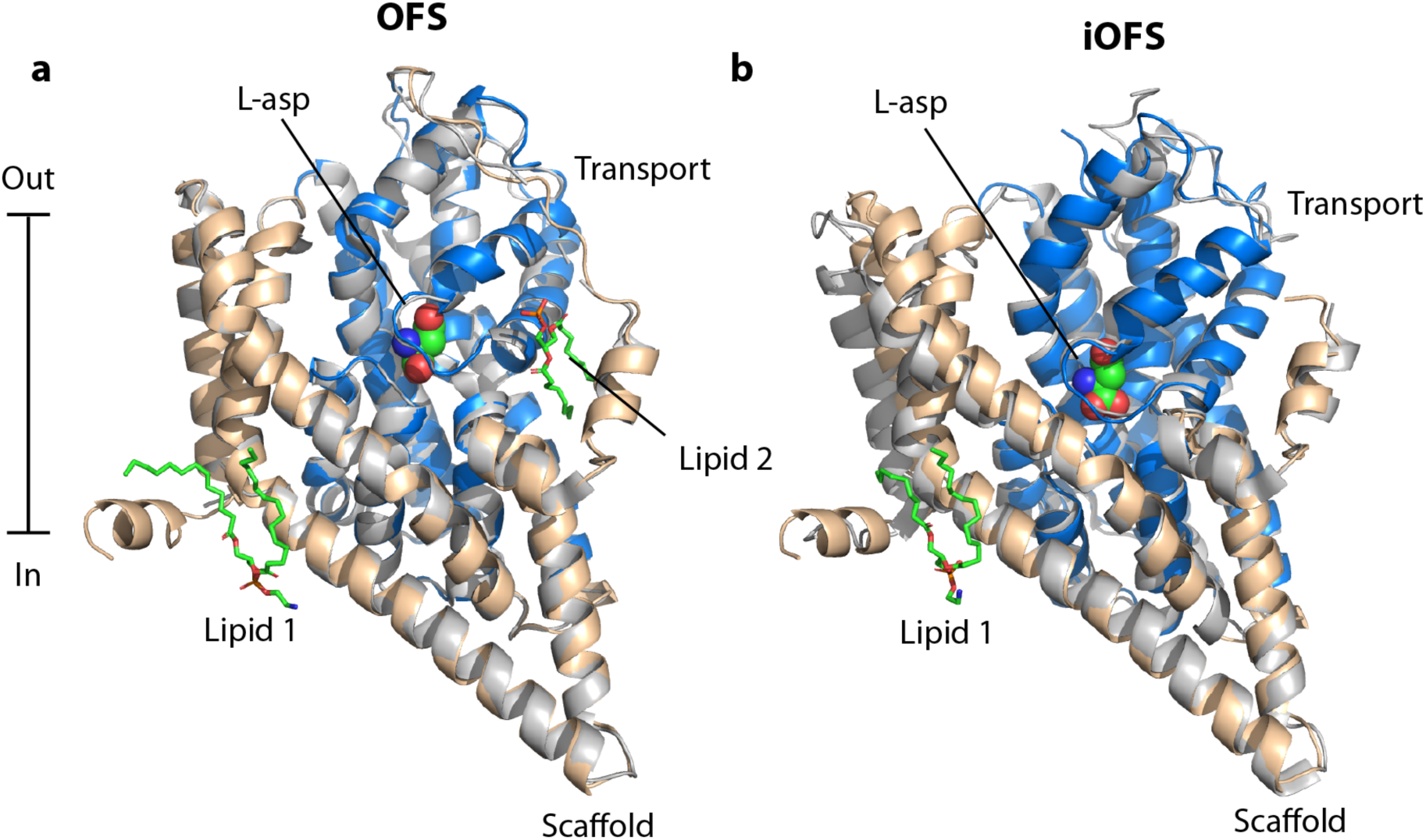
Structural comparison between Cryo-EM and crystal structures of GltPh in the OFS (a) and iOFS (b) conformations. Single protomers are viewed in the plain of the membrane and are depicted in cartoon representation. The crystal structures are colored gray, and the Cryo-EM structures are in color with scaffold domain wheat and the transport domain blue. Bound L-asp is shown as spheres and is colored by atom type. Two structurally well-defined lipid molecules were observed in the Cryo-EM maps of GltPh reconstituted into nanodiscs and visualized in OFS conformation. They are shown in stick representation, colored by atom type, and labeled Lipid 1 and 2. Lipid 1 was also observed in the maps of GltPh in iOFS conformation, while Lipid 2 is displaced by the shifted transport domain. Small differences in the conformation of the scaffold domain are observed between the iOFS seen in the crystal and in Cryo-EM structures.

**Figure S5.**
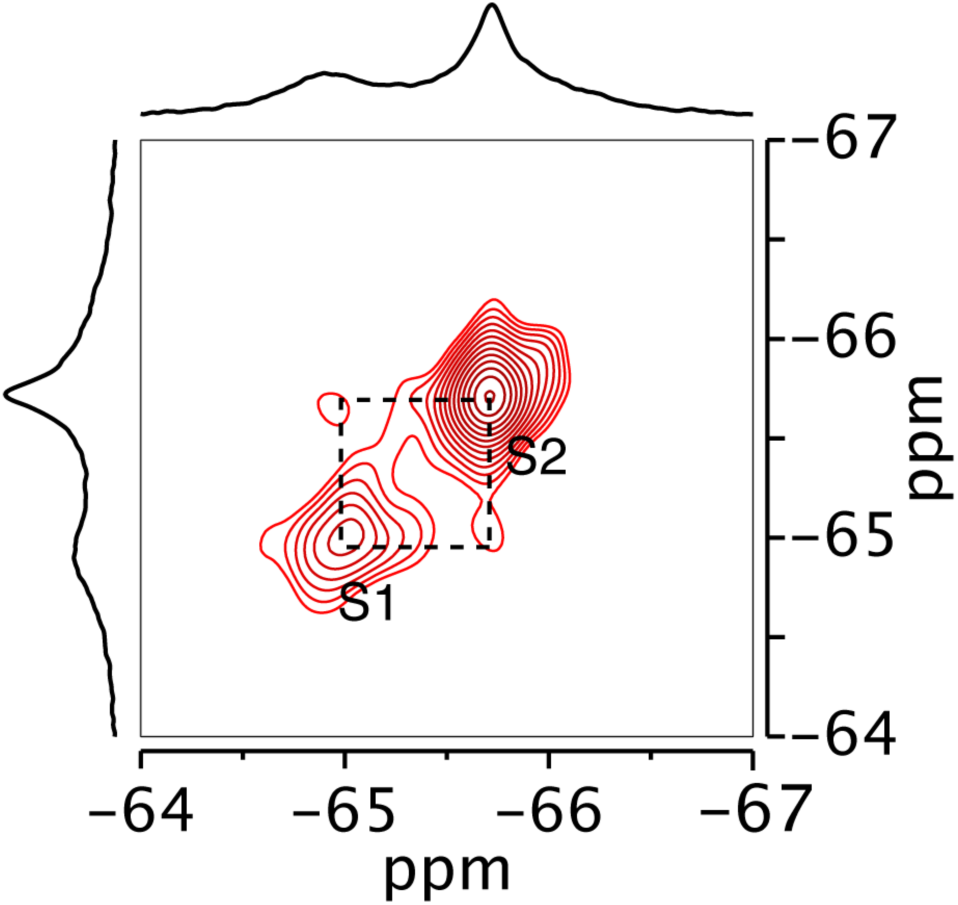
2D ^19^F EXSY spectrum of dHis/M385C-TET GltPh. Spectrum was recorded with mixing time of 0.4 s in the presence of 200 mM Na^+^ and 10 µM L-asp at 298K.

**Figure S6.**
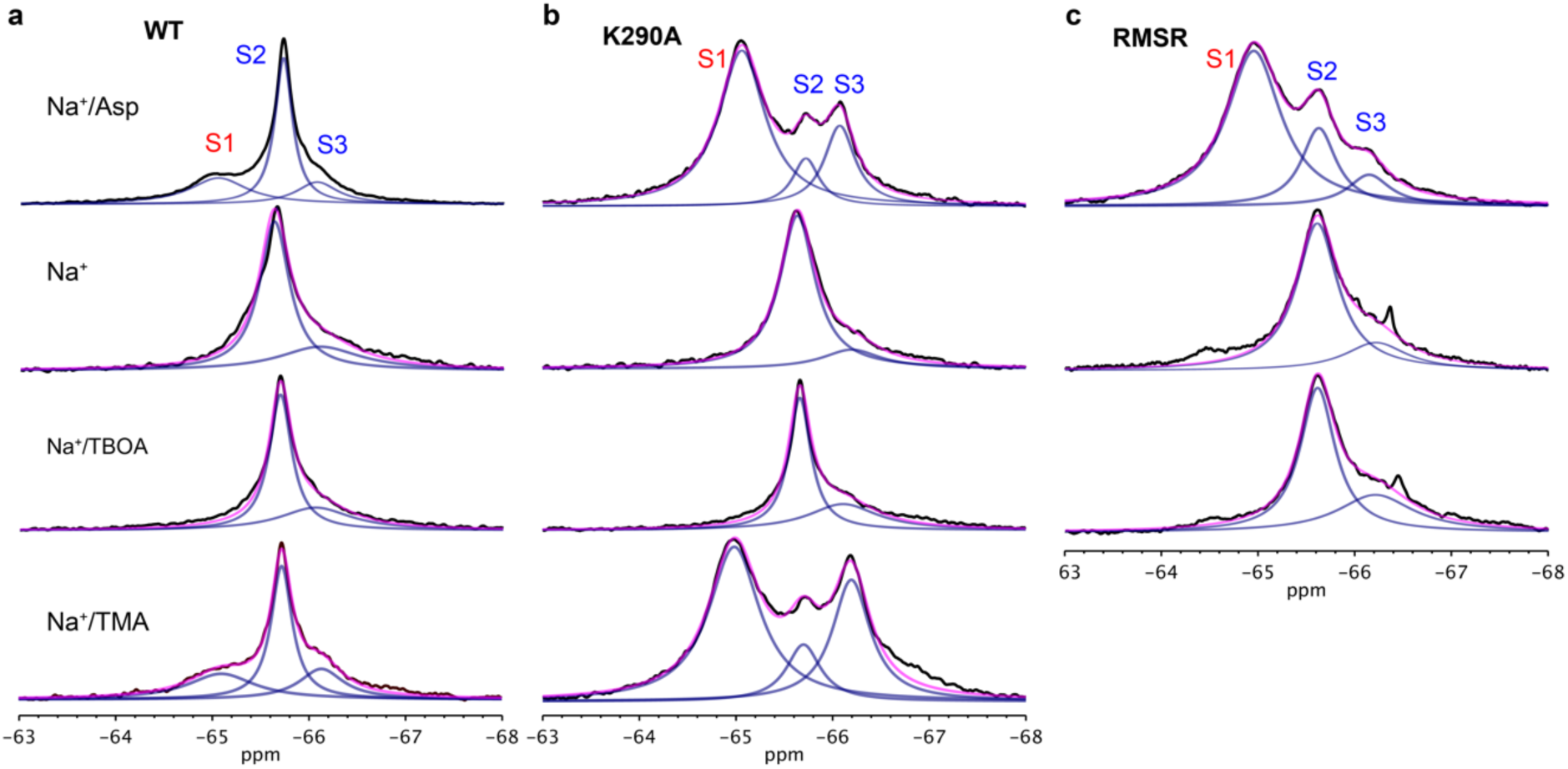
^19^F-NMR spectra of dHis/M385C-TET GltPh variants under different ligand bound conditions. (**a**), wild type (WT), (**b**), K290A, (**c**), RSMR. Experimental conditions from top to bottom are: 200 mM Na^+^ and 10 µM L-asp, 0.6 M Na^+^ only, 200 mM Na^+^ and 1 mM TBOA and 200 mM Na^+^ and 1.2 eq. TMA, respectively. All spectra were recorded at 293K. The spectra were deconvoluted into Lorentzian peaks S1, S2 and S3.

**Table S1.**
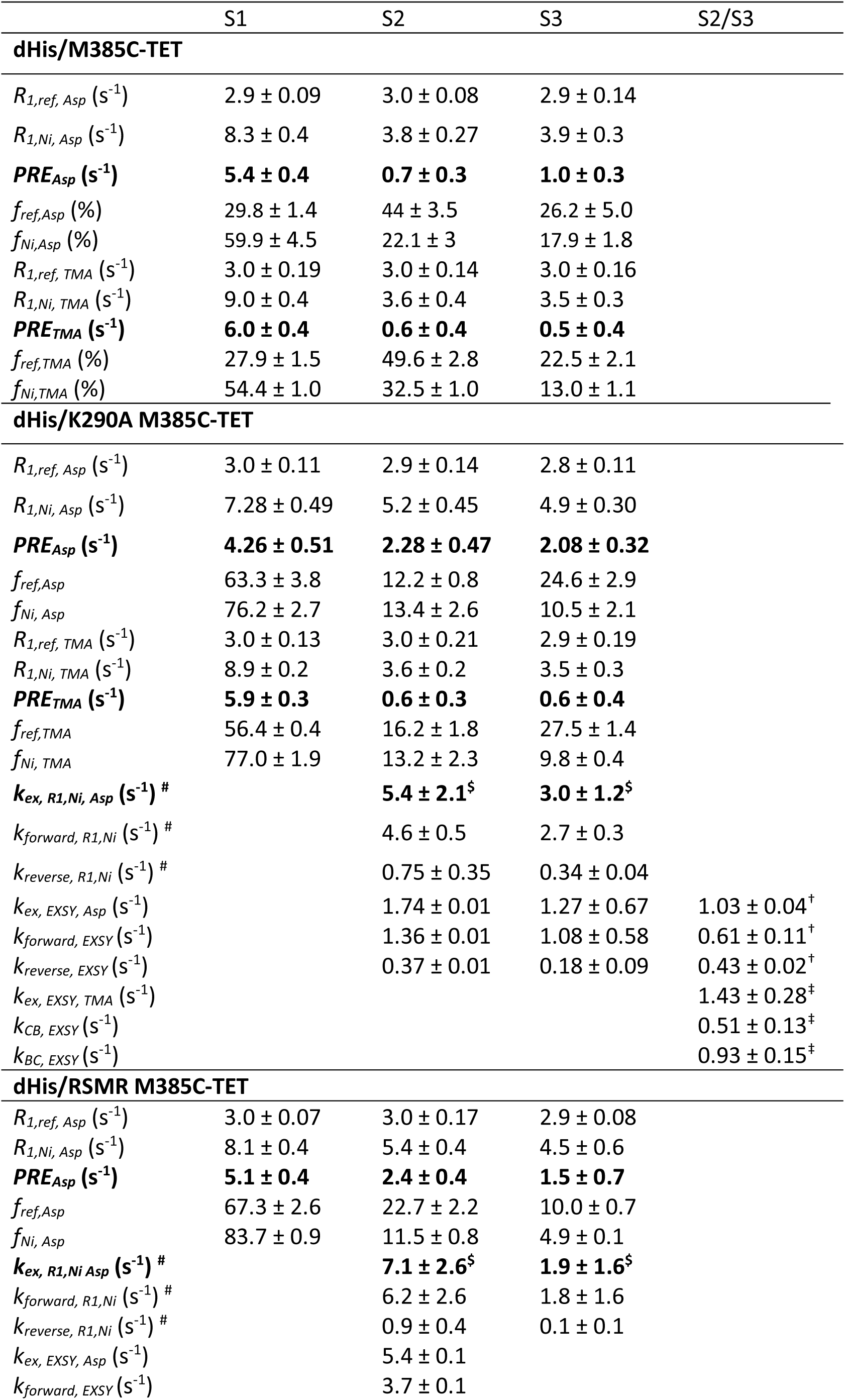
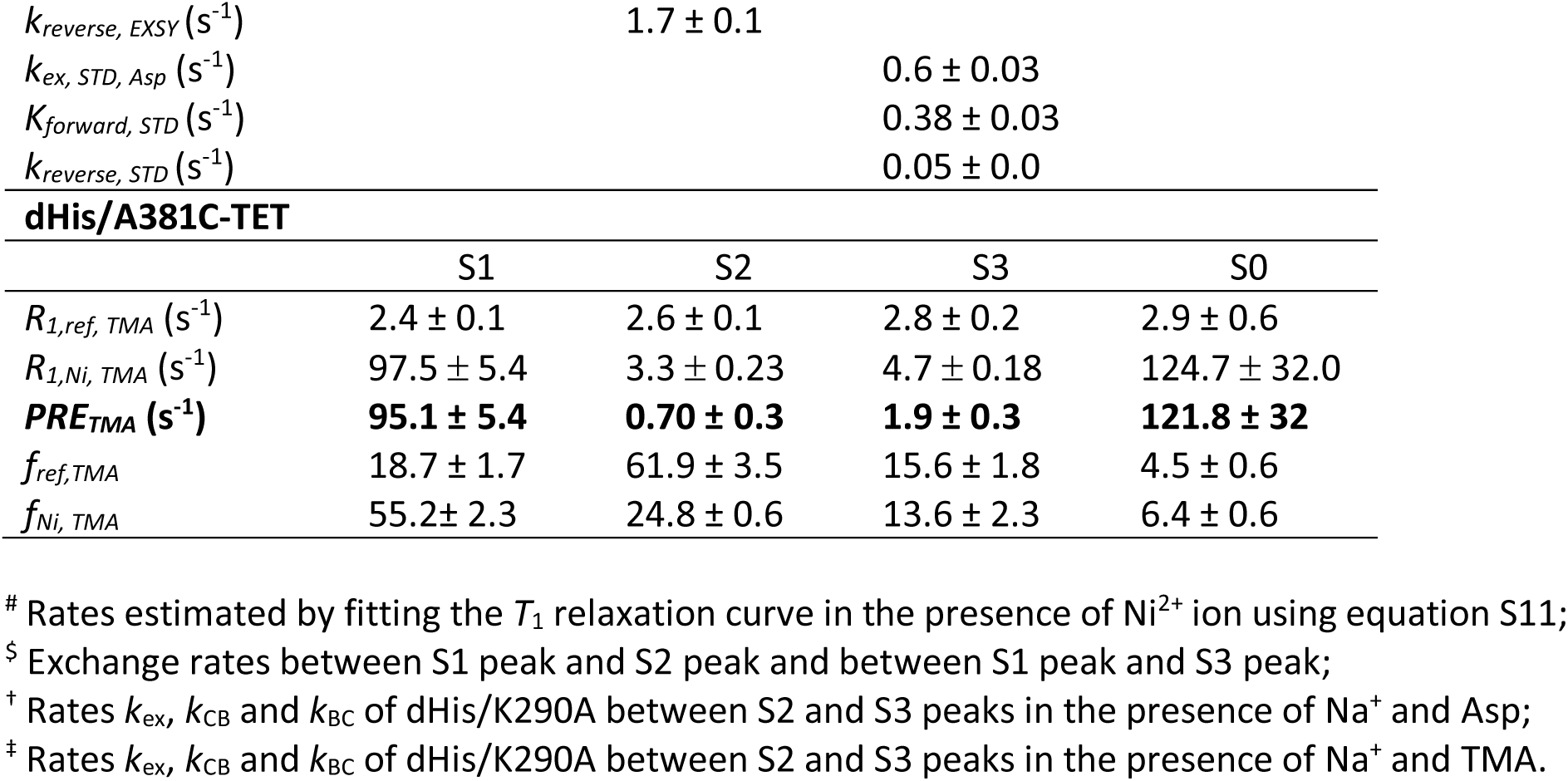
*R*_1_ relaxation rates and conformational exchange rates for GltPh variants.

**Table S2.** Cryo-EM data collection, reconstruction, and model refinement statistics Data collection

| **Data collection** |  |  |
| --- | --- | --- |
| Magnification | 81,000 |  |
| Exposure (e<sup>-</sup>/Å<sup>2</sup>) | 50.1615 |  |
| Defocus range (μm) | -1.5 to -2.5 |  |
| Pixel size (Å) | 0.53 |  |
| Micrographs (no) | 5236 |  |
| Initial particles (no) | 2,694,050 |  |
| Final particles (no) | 342,356 |  |
| **Reconstruction** | GltPh OFS | GltPh iOFS |
| Symmetry | C1 | C1 |
| Map resolution (Å) | 3.08 | 3.62 |
| FSC threshold | 0.143 | 0.143 |
| **Refinement** |  |  |
| **Initial model (PDB code)** | 2NWX | 3V8G |
| Map sharpening B-factor (Å<sup>2</sup>) | -104.1 | -146.6 |
| **Model composition** |  |  |
| Protein residues | 417 | 402 |
| Protein atoms | 3,178 | 3,029 |
| **Ligands** | 3 | 2 |
| **R.m.s. deviations** |  |  |
| Bond lengths (Å) | 0.005 | 0.007 |
| Bond angles (°) | 0.842 | 1.044 |
| **Ramachandran plot** |  |  |
| Favored (%) | 96.6 | 92.2 |
| Allowed (%) | 3.2 | 7.6 |
| Disallowed (%) | 0 | 0.25 |
| **Validation** |  |  |
| MolProbity score | 1.22 | 1.77 |
| Clashscore | 2.44 | 5.44 |
| Poor rotomers (%) | 0 | 0 |
| EMRinger score | 2.48 | 1.71 |

